# Transcriptome profile of Carrizo citrange roots in response to *Phytophthora parasitica* infection

**DOI:** 10.1101/598250

**Authors:** Zunaira Afzal Naveed, Jose C. Huguet-Tapia, Gul Shad Ali

## Abstract

*Phytophthora parasitica* is one of the most widespread *Phytophthora* species, which is known to cause root rot, foot rot/gummosis and brown rot of fruits in citrus. In this study, we have analyzed the transcriptome of a commonly used citrus rootstock Carrizo citrange in response to *P. parasitica* infection using the RNA-seq technology. In total, we have identified 6692 differentially expressed transcripts (DETs) among *P. parasitica*-inoculated and mock-treated roots. Of these, 3960 genes were differentially expressed at 24 hours post inoculation and 5521 genes were differentially expressed at 48 hours post inoculation. Gene ontology analysis of DETs suggested substantial transcriptional reprogramming of diverse cellular processes particularly the biotic stress response pathways in Carrizo citrange roots. Many *R* genes, transcription factors, and several other genes putatively involved in plant immunity were differentially modulated in citrus roots in response to *P. parasitica* infection. Analysis reported here lays out a strong foundation for future studies aimed at improving resistance of citrus rootstocks to *P. parasitica*.

## Introduction

Citrus is one of the most important commercial fruit crops, grown in more than hundred countries in tropical and subtropical areas (Talon and Gmitter, 2008). Altogether, citrus production is adversely impacted by numerous diseases, but major yield-affecting citrus diseases are distinct among different geographical regions of the world depending upon the climate, management practices and type of scion and root stocks (Davis, 1982). The *Phytophtora* disease complex on citrus is globally known to be the most damaging soil-borne disease of citrus. *Phytophthora parasitica, P. palmivora* and *P. citrophthora* are the three major *Phytophthora* species that can cause root rot, foot rot and sometimes in severe cases gummosis and brown rot of fruit (Savita and Nagpal, 2012). *P. parasitica* Dastur, also known as *P. nicotianae,* is the most widespread citrus infecting species throughout the tropical and subtropical areas of the world and is present in most citrus growing groves of Florida (Gade and Lad, 2018; Zitko and Timmer, 1994).

Phytophthora root rot is a serious problem of citrus particularly during nursery propagation of susceptible rootstocks, causing root decay and death of seedlings. Root rot in bearing orchards can cause slow tree decline and yield losses (Savita and Nagpal, 2012). Swingle citrumelo and Carrizo citrange are considered *Phytophthora* resistant rootstocks (Graham, 1995; Hutchison, 1974). In general, these rootstocks offer considerable resistance to *Phytophthora* diseases, but this resistance is usually broken down under disease complex situations when insect and multiple pathogen infestations acts in synergism causing severe damage to citrus trees. For example, the *Phytophthora-Diaprepes* complex, in which damage of citrus roots by larvae of *Diaprepes* root weevil tends to aggravate Phytophthora root rot, renders even highly resistant rootstocks susceptible to *Phytophthora* diseases (Graham et al., 2003). Interaction between Phytophthora root rot and citrus greening (also known as Huang Long Bing, HLB), which is the most devastating citrus disease in Florida, has also been observed (Graham et al., 2007). *Phytophthora spp.* prefers to infect new roots and HLB favors new growth of fibrous roots thus making this disease complex devastating for citrus trees. Understanding molecular bases of citrus defense against *Phytophthora* and its relevance to other citrus pathogens is required for designing long-term resistance development strategies to combat losses caused by these noxious pathogens.

The first citrus disease caused by *P. parasitica* was reported back in 1832, well before the advent of the discipline of plant pathology. Being so ancient and a widespread problem, a lot of research had been conducted to understand the biology, etiology and control of this disease. However, little is known about the molecular basis of citrus-*Phytophthora* interactions. A foliar microarray based differential gene expression analysis among resistant and susceptible hybrids and their resistant (*Poncirus trifoliata* cv. Rubidoux) and susceptible (*Citrus sunki)* parents was conducted in response to stem inoculation of *P.parasitica* (Boava et al., 2011). Several genes that were differentially expressed during these comparisons were suggested to play a role in the resistance or susceptibility of these hosts to citrus gummosis. Recently, Dalio et al. reported the molecular basis of compatible and incompatible citrus-*P. parasitica* interaction by using susceptible (*Citrus sunki*) and resistant (*Poncirus trifoliata*) citrus rootstocks, respectively, by comparing the gene expression of defense related host genes through quantitative PCR. They have shown significant induction of key defense genes in susceptible rootstock whereas little or no changes in the expression level of tested defense genes in the resistant rootstocks were observed by them in response to pathogen (Dalio et al., 2017). Similarly, on the pathogen side, transcriptional profiling of *P. parasitica* during citrus gummosis was conducted to identify pathogenicity factors during infection (Rosa et al., 2007). These reports of citrus-*Phytophthora* interactions are either confined to the above ground infections where mostly leaf tissues were used for transcriptome analysis or restricted to few defense related genes in roots, but so far, no study has been done to determine whole transcriptional profiling of citrus roots during Phytophthora root rot.

Innate immune system of plants comprises of two major layers of defense responses (Dodds and Rathjen et al, 2010). In the first layer of defense, pattern recognition receptors (PRRs), which are localized to the host cell membranes, recognize highly conserved molecules or structures of pathogens known as pathogen associated molecular pattern (PAMP) and activate PAMP-Triggered Immunity (PTI); PTI leads to an array of basal defense responses that combat pathogen invasion. However, when the pathogen manages to survive PTI then the second layer of immunity called effector triggered immunity (ETI), which is faster and more robust than PTI, is activated. ETI is dependent on pathogen effectors, which are proteins secreted by pathogens inside host cells, where they target and modulate the activity of host proteins for inactivating defenses and procuring nutrients. In incompatible interactions, plant resistance (R) proteins perceive these modulations leading to the activation of downstream defense signaling, which ultimately result in local cell death or hypersensitive response (HR) to prevent the spread of the pathogen. Most of the *R* genes code for intracellular receptor proteins, which contains nucleotide binding site (NBS) and leucine rich repeat (LRR) domains (Jones and Dangl, 2006). However, some membrane associated *R* genes that function as PRRs have also been reported (Zipfel et al., 2006). Based on their functional domains, *R* genes are broadly categorized into five classes: (i). CNL (CC-NBS-LRR), which comprise of at least one N-terminal coiled-coil (CC) domain in addition to the typical NB-LRR domain, (ii). TNL (TIR-NBS-LRR), which contains an N-terminal Toll-Interleukin Receptor (TIR) like domain in addition to the NB-LRR domain, (iii). RLPs (Receptor-Like Protein), which consists of an extracellular LRR domain, which usually associates with a cytoplasmic serine threonine kinase, (iv). RLK (Receptor-Like Kinases), which contains an extracellular leucin rich repeat (LRR) and a cytoplasmic kinase domain, and (v). Others, which comprise of all other resistance genes without any defined conserved domains (Sanseverino et al., 2009). Several *P. infestans* resistance genes (*Rpi*) have been identified in tomato, potato and other plants and they are currently being used in many *Phytophthora* resistant cultivars (Rodewald and Trognitz, 2013; Zhu et al., 2012). But most of these *Rpi* genes are race-specific in recognizing their cognate *Phytophthora* effector counter parts (coded by avr genes) that are diverse not only among different *Phytophthora* species but also among different races and strains of same species (Rodewald and Trognitz, 2013). Two *R* genes, a TIR-NB-LRR and RSP4 were shown to be involved in resistance against *P. parasitica* in a citrus cultivar (*Poncirus trifoliata* cv. Rubidoux) (Boava et al., 2011). In addition to local cell death (HR), *R* genes are also known to activate prolonged resistance by inducing phytohormones and *pathogenicity related* genes (*PR* genes) that collectively give rise to broad spectrum systemic acquired resistance (SAR) against future infections (Jones and Dangl, 2006). SAR and Induced Systemic Resistance (ISR), which is incited by soil borne infections can prevent infections in aerial foliage (Park et al., 2007; Vernooij et al., 1994).

In this study, using the RNA-seq approach, we have analyzed whole transcriptome of Carrizo citrange rootstock in response to *P.parasitica* at 24 and 48 hours post inoculation (hpi) to determine the transcriptional changes during this tolerant citrus-*Phytophthora* interaction. Our major focus was to identify citrus genes potentially involved in defense particularly *R* genes. Findings of our analyses provide insights into how citrus roots provide resistance to *Phytophthora*, and these findings provide a strong basis to further explore the molecular basis of tolerance offered by Carrizo rootstock. Potential uses of the outcomes of our analyses for developing rootstocks that are tolerant to *Phytophthora*, especially in the context of disease complexes are discussed.

## Materials and methods

### *P. parasitica* inoculation and sample collection and sequencing

*P. parasitica* strain 13-723A was isolated from citrus plants grown in local nursery in Florida. It was grown on 10% solid V8 medium plates at 12:12 hours dark:light cycle at 25 °C for 10 days. Zoospores were induced by applying dehydration stress on cultures from day 5 to day 7. On 10^th^ day of culture, plates were flooded with sterilized water to induce release of zoospores. Zoospores were filtered through two to four layers of cheesecloth and adjusted to 10^5^ cells/ml. Roots of sixty days old seedlings of Carrizo citrange rootstock grown under controlled conditions in sterilized soil were inoculated with 50 ml zoospore suspension. Mock inoculation was done using 50 ml sterilized water without zoospores.

Root samples from pathogen and mock treatments were collected at 24 hours post inoculation (hpi) and 48 hpi. Roots were rinsed in sterile water, immediately frozen in liquid nitrogen and stored at −80 °C until further use. Total RNA was extracted from frozen root tissues and six cDNA libraries were constructed and sequenced using the paired-end (PE) 125 x 2 sequencing reactions using the Illumina HiSeq 2500 platform.

### RNA-Seq quality assessment

All raw reads were assessed for quality using FastQC (v0.11.5) and the CLC Genomics Workbench 9.0 (Qiagen). All PE raw reads were processed for trimming the Illumina sequencing adaptors and low-quality reads with the Trim Galore software (version 0.4.3). Quality and trim reports are provided in Appendix S1.

### Reference genome mapping

Clean reads, after removing adaptors and low-quality reads, from all samples were mapped to two *Citrus sinensis* genomes (Ridge Pineapple and Valencia) and to one Swingle citrumelo (*Citrus x paradisi x Citrus trifoliata*) genome separately. Samples containing *Phytophthora parasitica* reads were mapped to the *P. parasitica* INRA-310 genome (BROAD institute). Reference genome mapping was done using the bowtie2 algorithm of tophat aligner (Trapnell et al., 2009; Langmead et al., 2009).

### *De novo* assembly and differential expression analysis

Pathogen contamination in the *P.parasitica* treated samples libraries were removed by filtering out the reads mapped to *P.parasitica* reference genome. Clean reads from all seven samples were *de novo* assembled using Trinity (Grabherr et al., 2011) to get a pooled *de novo* reference assembly for citrus. Although pathogen reads were filtered before generating pooled host assembly, pathogen contamination were still there. In order to remove that, a local BLAST search was done against two customized databases based on all available Citrus and *Phytophthora* spp. genomes (Table S2) and all putative pathogen transcripts were removed.

Assembly optimization was done by following the previously described approach with few amendments (McCann et al., 2017). Briefly, all Trinity assembled transcripts were fed to TransDecoder package (Haas et al., 2013) for the prediction of open reading frames (ORFs). Longest ORF peptide sequences were then subjected to BLASTp against uniport protein database as well as scanned for protein domains using Pfam. Results from BLASTp and Pfam were integrated to retain the coding transcripts and rest were all filtered out. Isoforms having more than 98% similarities were removed to keep only the unique coding transcripts.

Optimized assembly was used as a reference for further analysis of citrus transcriptome and cleaned paired end reads from all samples were subjected to transcript quantification by using RSEM (RNA-seq by Expectation Maximization) program (Li and Dewey, 2011), which is incorporated in the Trinity pipeline. Abundance estimation steps of RSEM involve mapping of clean PE reads of each sample back to the *de novo* pooled assembly followed by calculation of gene expression by counting reads mapped to trinity genes and transcripts. Read counts from all samples were combined in a matrix and normalization was done using the TMM method of RSEM (Li and Dewey, 2011). Quality of pooled assembly was checked by calculating N50, ExN50 statistics (Figure S1) and other summary statistics (Table 1). Differential expression analysis among the mock and *P. parasitica*-infected treatments at each time point was done using the EdgeR package (Robinson et al., 2010; Seyednasrollah et al., 2013). Since we had two replicates of the pathogen-treated samples and one replicate for the mock-inoculated samples at 24 hpi and 48 hpi, and one replicate at the 0 hr control, the dispersion parameter in EdgeR was set to 0.1. Differential expression was analysis was done based on both coding isoforms count and unigene count. Whole analysis pipeline is summarized in Figure S2.

**Table 1.**
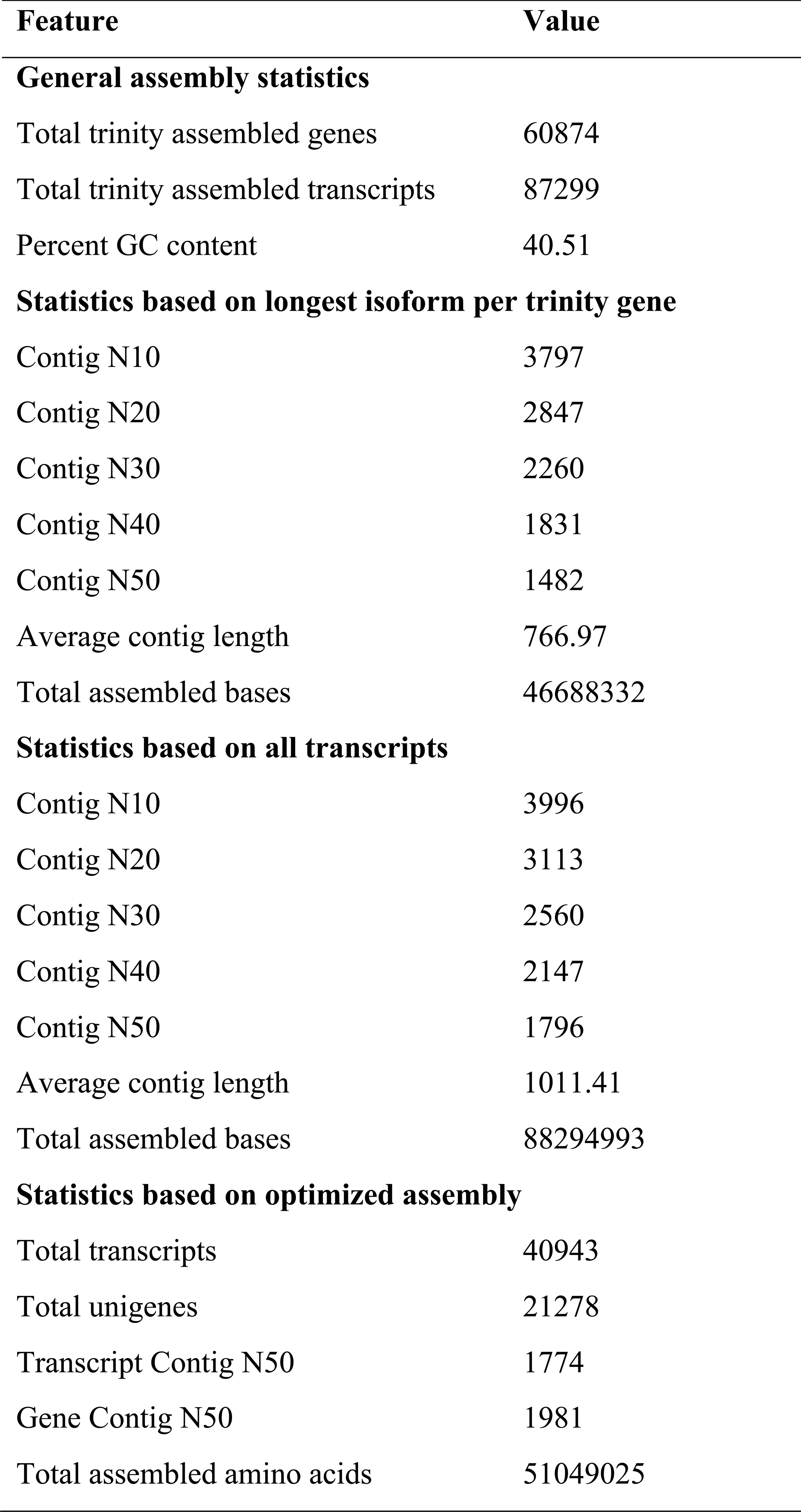
The statistics of Trinity generated *de novo* assembly.

| **Feature** | **Value** |
| --- | --- |
| **General assembly statistics** |  |
| Total trinity assembled genes | 60874 |
| Total trinity assembled transcripts | 87299 |
| Percent GC content | 40.51 |
| **Statistics based on longest isoform per trinity gene** |  |
| Contig N10 | 3797 |
| Contig N20 | 2847 |
| Contig N30 | 2260 |
| Contig N40 | 1831 |
| Contig N50 | 1482 |
| Average contig length | 766.97 |
| Total assembled bases | 46688332 |
| **Statistics based on all transcripts** |  |
| Contig N10 | 3996 |
| Contig N20 | 3113 |
| Contig N30 | 2560 |
| Contig N40 | 2147 |
| Contig N50 | 1796 |
| Average contig length | 1011.41 |
| Total assembled bases | 88294993 |
| **Statistics based on optimized assembly** |  |
| Total transcripts | 40943 |
| Total unigenes | 21278 |
| Transcript Contig N50 | 1774 |
| Gene Contig N50 | 1981 |
| Total assembled amino acids | 51049025 |

### Functional annotation of transcripts

Differentially expressed transcripts were annotated using Blast2GO with the default parameters (Conesa et al., 2005). CloudBlast search option was used to BLAST DETs against *viridiplantae* database with an expect value (e-value) of 1.0e^-3^. To identify *R*-genes, a local BLASTx search of optimized assembly transcript sequences against all available plant *R* gene protein sequences in the Plant Resistance Gene database (PRGdb) was performed with an e-value 1.0E^-10^ and *R* genes were selected if percent identity was greater than 80 and bit score was greater than 100 (Sanseverino et al., 2012).

For GO term enrichment analysis, GO terms for DETs obtained from annotation results were fed to the Singular Enrichment Analysis (SEA) tool of AgriGO using customized parameters (Du et al., 2010). Reduce and visualize gene ontology (REViGO) was used to summarize and visualize most significantly enriched GO terms (Supek et al., 2011).

KEGG (Kyoto Encyclopedia of Genes and Genomes) enzyme codes for DETs were obtained through Blast2GO plugin. In addition to that all DETs were fed to KAAS (KEGG automatic annotation server) to assign KO terms (Moriya et al., 2007) and Pathview was used to generate KEGG pathways (Luo et al., 2017). Mapman software was also utilized to view DETs in biological processes (Thimm et al., 2004; Usadel et al., 2009). Mercator web application (Lohse et al., 2013) and homology search for citrus orthologs were used to assign MapMan bins.

### Validation of DETs by Quantitative Real time PCR

To validate differential expression results obtained by following Trinity-RSEM-EdgeR pipeline, ten DETs were subjected to reverse transcriptase PCR (RT-PCR) for cDNA synthesis followed by quantitative real time PCR (qRT-PCR) by following the previously published protocol (El-Sayed et al., 2015; Patel et al., 2015). GADPH gene was used as a reference to calculate the relative expression values. Gene specific primer pairs were designed using NCBI primer BLAST tool (https://www.ncbi.nlm.nih.gov/tools/primer-blast/). Primers used for this experiment are given in Table S3. Melting curve analysis was done to ensure amplification of single gene product by each primer pair.

## Results

### *Phytophthora parasitica* infects and colonize Carrizo citrange roots

Carrizo citrange has been reported to be tolerant to *P.parasitica* infection. To investigate whether *P.parasitica* infects Carrizo citrange roots or not, we inoculated roots of 60 days old plants with zoospores and followed infection dynamics studies under confocal microscope. Zoospores attachment, encystement and cyst germination on roots surface were observed at approximately 3 hpi (Figure 1A). Inter and intracellular hyphal growth was clearly observed at 24 hpi followed by heavy colonization at 48 hpi. Haustoria like structures were also clearly observed at 24 and 48 hpi (Figure 1B and C). Contrastingly, no pathogen structures were observed during *P. parasitica* interaction with *P. trifoliata* that is another resistant citrus rootstock (Dalio et al., 2017), suggesting that different rootstocks respond differently to different strains of *P. parasitica*. Like all other *Phytophthora* species, *P.parasitica* is a hemi-biotrophic pathogen. Infection studies of *P.parasitica* in *A. thaliana* roots have revealed a short biotrophic phase (between 0 to 24 hpi) followed by quick switch to a necrotrophic phase at about 30 hpi (Attard et al., 2010). Although, *P.parasitica* infection and colonization was observed on Carrizo citrange roots, disease symptoms in aerial parts of the plants were very mild. To capture transcriptional changes in the infected roots and understand molecular basis of disease resistance, we collected infected root samples at 24 hpi and 48 hpi for RNA-seq analyses.

**Figure 1.**
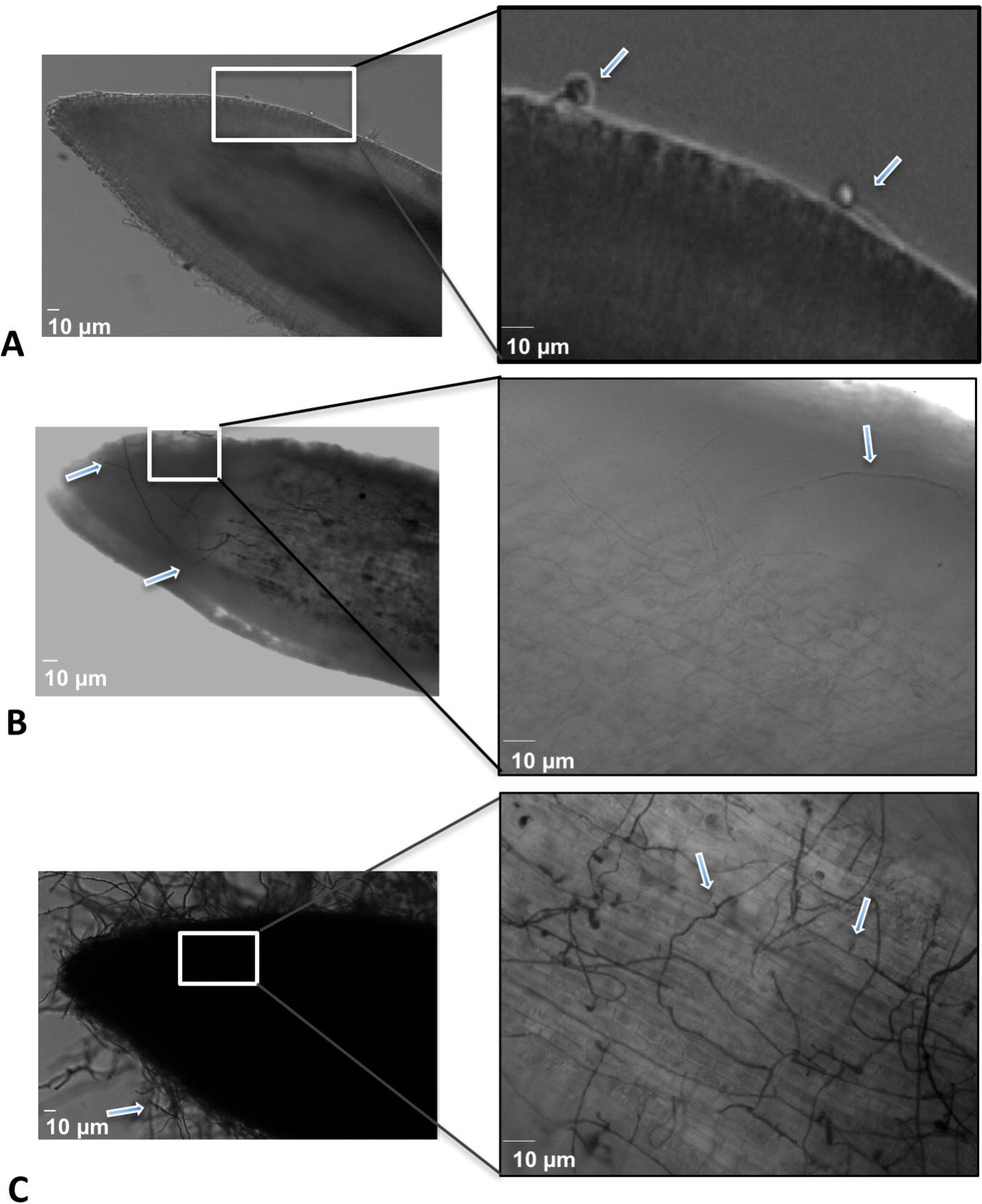
Infection dynamics of *Phytophthora parasitica* in Carrizo roots from 3hpi to 48hpi. *P. parasitica* zoospore attachment and germination on the root surface indicated by arrows, observed at 3hpi in confocal microscope under Differential interference contrast white field (DICWF) filter (A). Inter and intracellular hyphal growth (indicated by arrow) was observed at 24hpi (B). A heavy colonization of *P. parasitica* mycelium was seen at 48hpi. After trypan blue staining hook like invaginations that are probably haustoria are clearly visible, indicated by arrows (C).

### mRNA-seq data analyses and quality assessment

Total RNA was isolated from infected and mock-treated roots, processed and sequenced using paired end (PE) 125 x 2 reaction on the Illumina HiSeq 2500 platform according to the manufacturer’s protocol. In total, 165 million PE reads (20 billion base pairs) with an average length of 120 bp were obtained. Average Phred scores of all reads was above 30 and more than 70% reads had Phred score > 35, indicating very high-quality sequence data (Appendix S1). After removing Illumina adaptors and low-quality reads, approximately 165 million clean reads comprised of 19.8 billion base pairs remained, providing us with sufficient depth for differential expression analyses.

### Reference Genome Mapping

All four *P. parasitica* containing samples were mapped to the *P. parasitica* INRA-310 genome. At 24hpi both replications of pathogen treated citrus roots showed 0.8% mapping coverage on *P. parasitica* genome. Whereas, a higher mapping coverage of 1.8% was observed at 48 hpi in both samples (Table S1), indicating increased growth of *P. parasitica* on citrus roots from 24 to 48 hpi. Carrizo citrange is a hybrid of *Citrus sinensis* and *Poncirus trifoliata*. Currently, two *C. sinensis* genomes are available but none for *P. trifoliate* (Adhikari et al., 2012; Wang et al., 2014). However, draft genome of Swingle citrumelo, which is also a hybrid of *C. paradisi Macf* and *P. trifoliata*, is available (Zhang et al., 2016) and it could be used as a reference to map the Carrizo citrange reads that correspond to the *P. trifoliata* genome. To determine the proportions of these two, one parent (*C. sinensis*) and one step sibling (Swingle) genomes in Carrizo transcriptome, all read samples were separately mapped to these three reference genomes. Results of mapping to all three genomes showed only a 55-73 % coverage of the Carrizo transcriptome suggesting that a significant proportion of the Carrizo genome might be missing from these three genomes (Table S1). Therefore, to improve coverage, we constructed a *de novo* genome assembly and used it as a reference for differential gene expression analyses as follows.

### *De novo* transcriptome assembly and gene expression analysis

After reference genome mapping, reads mapped to *P.parasitca* genomes were filtered out and pathogen free clean reads from all seven samples were used to generate a pooled *de novo* transcriptome assembly using Trinity (Grabherr et al., 2011), one of the most widely used *de novo* transcriptome assembler for short reads in both animal (Hsu et al., 2017; McCann et al., 2017) and plant transcriptome studies (Evangelisti et al., 2017; Guo et al., 2016; Xiong et al., 2017; Yang et al., 2017). Altogether, 60874 trinity genes and 87299 trinity assembled transcripts were obtained with an N50 values of 1428 bp and 1796 bp, respectively (Table 1). In additional checks for contamination, 1043 *Phytophthora* genes (comprised of 1082 trinity transcripts) were removed.

Trinity transcripts are artificially assembled so chances of redundant transcripts are high. Thus, before moving further an optimization of Trinity constructed de novo assembly is necessary to get rid of false positives and to save statistical power in the downstream analysis (McCann et al., 2017; Ono et al., 2015). Assembly optimization was done by running homology-based search using BLAST and Pfam on Transdecoder predicted ORFs (see methods for detail) and 21,278 coding unigenes comprised of 40,943 coding transcripts were obtained (Table 1).

To estimate transcript abundance, reads were mapped back to the optimized reference assembly and gene expression in terms of normalized FPKM (fragment per kilo base of exon per million fragments mapped) was calculated using the TMM method in RSEM. RSEM is a Trinity supported program for accurate gene expression quantification using *de novo* assembled reference transcriptomes (Haas et al., 2013; Li and Dewey, 2011). To check the quality of our reference assembly, ExN50 statistics that refers to N50 of the top most highly expressed transcripts was computed from normalized expression data, resulting in E90N50 value of 2056 bp, which means that 90% of the top highly expressed transcripts have an N50 value of 2056 bp and indicates a very high-quality assembly (Figure S1).

### Estimation of variation among samples

To determine the reproducibility of biological replicates and variations among *P. parasitica*-infected and mock-inoculated samples over both time points, we performed correlation and principal components analysis (PCA) among all samples. Sample correlation matrix based on log2 transformed counts revealed that pathogen-treated replicates and mock-treated controls at both time points displayed high correlation with each other. As is shown in the dendogram in Figure 2A, biological replicates at each time point clustered together indicating high reproducibility of the data. Consistent with the expectations of the experimental conditions, *P. parasitica-*infected samples at both time-points also clustered together. Similarly, mock-treated controls at both time points also clustered together but separate from the cluster of *P. parasitica-*infected samples. These results were validated with PCA analysis (Figure 2B). PCA plots of PC1 vs PC2 and PC2 vs PC3 revealed very little variation among replicates at both time points (Figure 2B). Similarly, mock-inoculated samples both at 24 hpi and 48 hpi clustered together. As expected, PC1 (42.9%) explained most of the variations among mock-inoculated and pathogen-infected root samples, whereas, PC2 (18.7%) explained variation among the mock-treated and the 24 hpi and 48 hpi samples (Figure 2).

**Figure 2.**
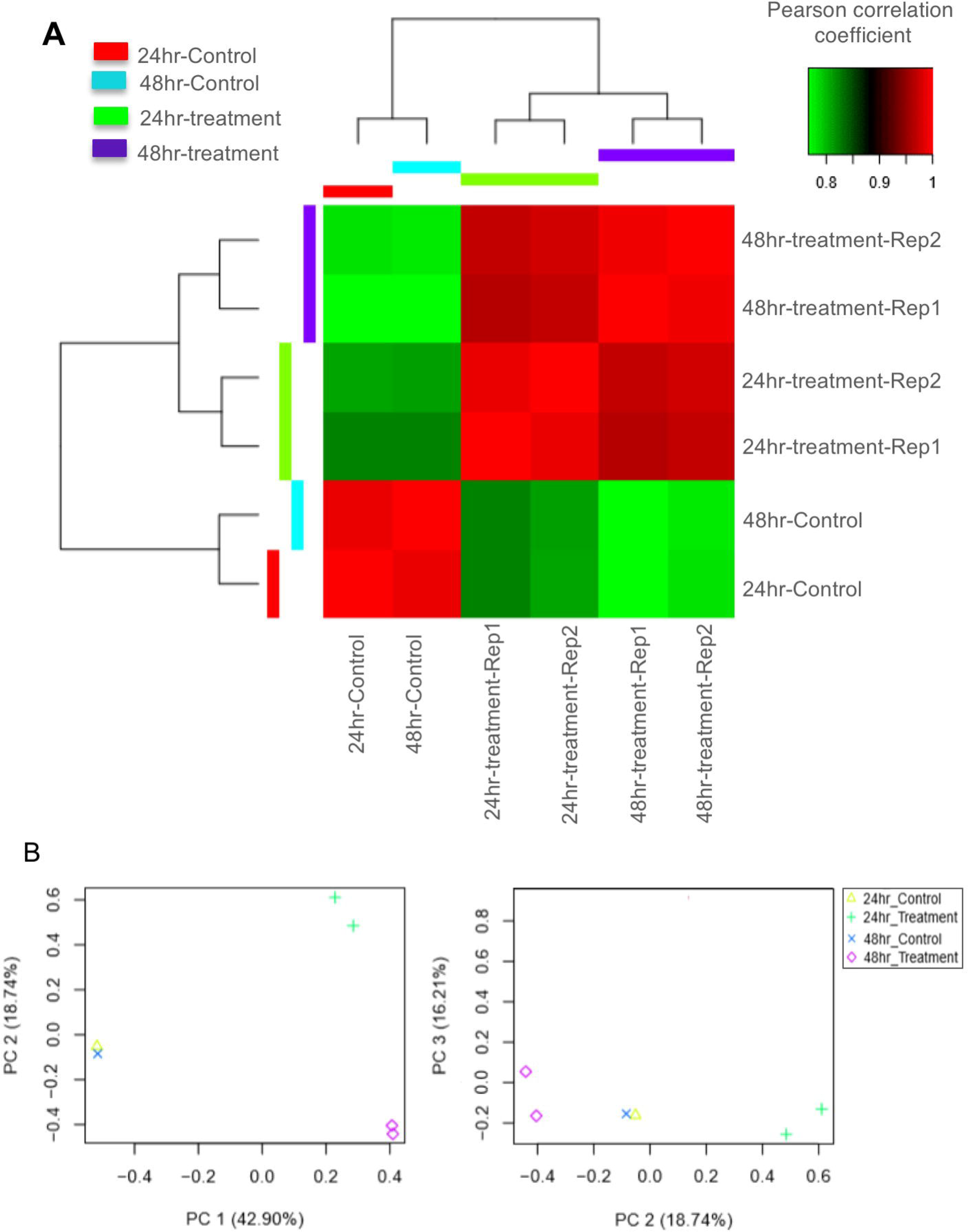
Sample correlation and Principle component analysis (PCA). Dendogram and sample correlation matrix heatmap shows high correlation between biological replicates than among treatments. Green to red coloring refers to low to high correlations (A). PCA analysis shows that biological replicates of *P. parasitica*-inoculated samples at both time points (2hr_Treatment and 48hr_Treatment) cluster together reassuring the heatmap analyses shown in A. Similarly, mock-treated samples at both time points also clustered together (B).

### Differential expression analysis of citrus genes in response to *P. parasitica* infection

Differential expression analysis was performed using EdgeR package, which has been shown to work better for data with fewer replicates (Robinson et al., 2010; Seyednasrollah et al., 2013). Volcano plots of all pairwise comparisons presenting log fold change distribution of differentially expressed transcripts along the *x*-axis and False Discovery Rate (FDR) values on the *y*-axis showed differential expression of numerous transcripts among all comparisons (Figure S3).

Differentially expressed transcripts (DETs) with log2 fold change >2 threshold and FDR < 0.005 were selected for further analysis. Overall hierarchical cluster heatmap is shown in Figure 3A. Hierarchical cluster analysis of DETs across all samples revealed four sub-clusters representing distinct differential expression patterns among infected and mock inoculated roots (Figure 3B). Sub-cluster 1 consisted of 2290 down-regulated DETs that are further down-regulated from 24 hpi to 48 hpi, sub-cluster 2 contained of 1924 up-regulated DETs that are slightly up-regulated from 24 hpi to 48 hpi and sub-cluster 3, comprised of 419 transcripts that are very highly up-regulated compared to controls and continue to increase expression from 24 to 48 hpi. Whereas, sub-cluster4 represents down-regulated DETs which have same expression at both time points (Figure 3B).

**Figure 3.**
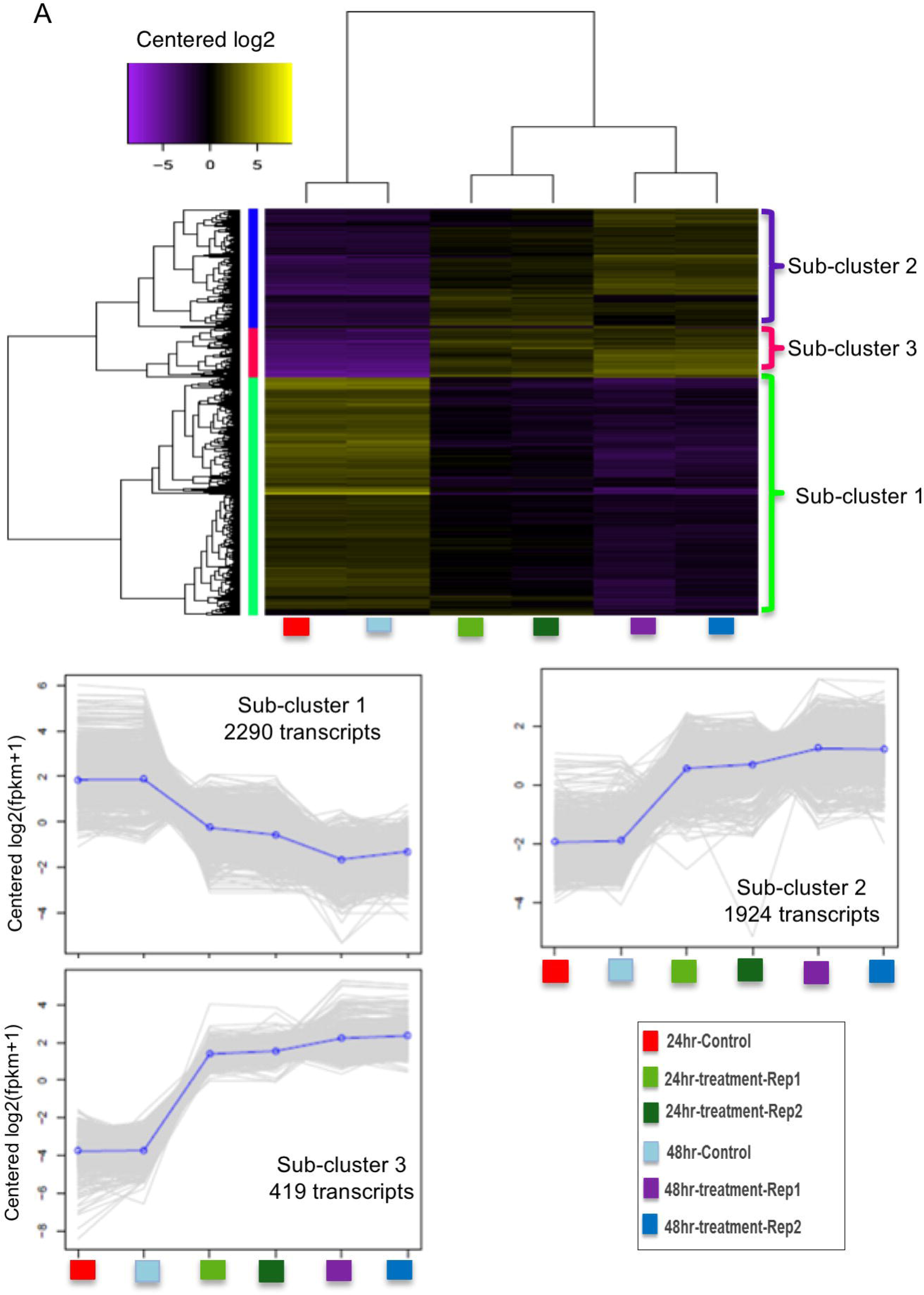
Hierarchical cluster analysis of differential expressed transcripts (DETs) across all samples. Heatmap showing overall hierarchical clustering of all DETs in citrus roots in response to *P. parasitica* infection. Yellow color is indicating induced expression and purple is pointing towards repression (A). Based on similarities in their expression profiles, total DETs are divided in three different sub-clusters. Sub-cluster 1 is showing highest number of DETs with increased expression from mock vs. *P. parasitica* treated samples at both time points. Similarly, second sub-cluster is representing down-regulated DETs upon infection at both time points. Third cluster is not only representing highly up-regulated DETs in mock vs infection but also slight expression variations from 24hpi to 48hpi (B).

In the 24 hpi treatment vs. 24 hpi mock comparison, a total 3960 DETs were identified with 2193 upregulated and 1767 downregulated. In the 48 hpi treatment vs. 48 hpi mock comparison, we identified 5521 DETs with 2526 upregulated and 2995 downregulated. Whereas, only 690 DETs were observed in the 24 hpi and 48 hpi pathogen-infected comparison. Venn diagram analysis of DETs among all pairwise comparisons revealed 844 and 2175 DETs uniquely expressed in citrus roots in response to *P. parasitica* at 24 hpi and 48 hpi respectively (Figure S4).

### Functional annotation of citrus DETs and mapping on plant defense pathways

Comprehensive functional annotation of DETs was done by using Blast2GO software (Conesa et al., 2005). Annotation statistics of DETs from all the databases is shown in Figure 4. Out of the total 6692 citrus DETs, BLAST hits were found for 6544 DETs. The top hits species distribution showed that majority of sequences were aligned to *Citrus sinensis* followed by *C. clementina* whereas few hits were found against other six *Citrus* spp. with yet available genome information (Figure S5).

**Figure 4.**
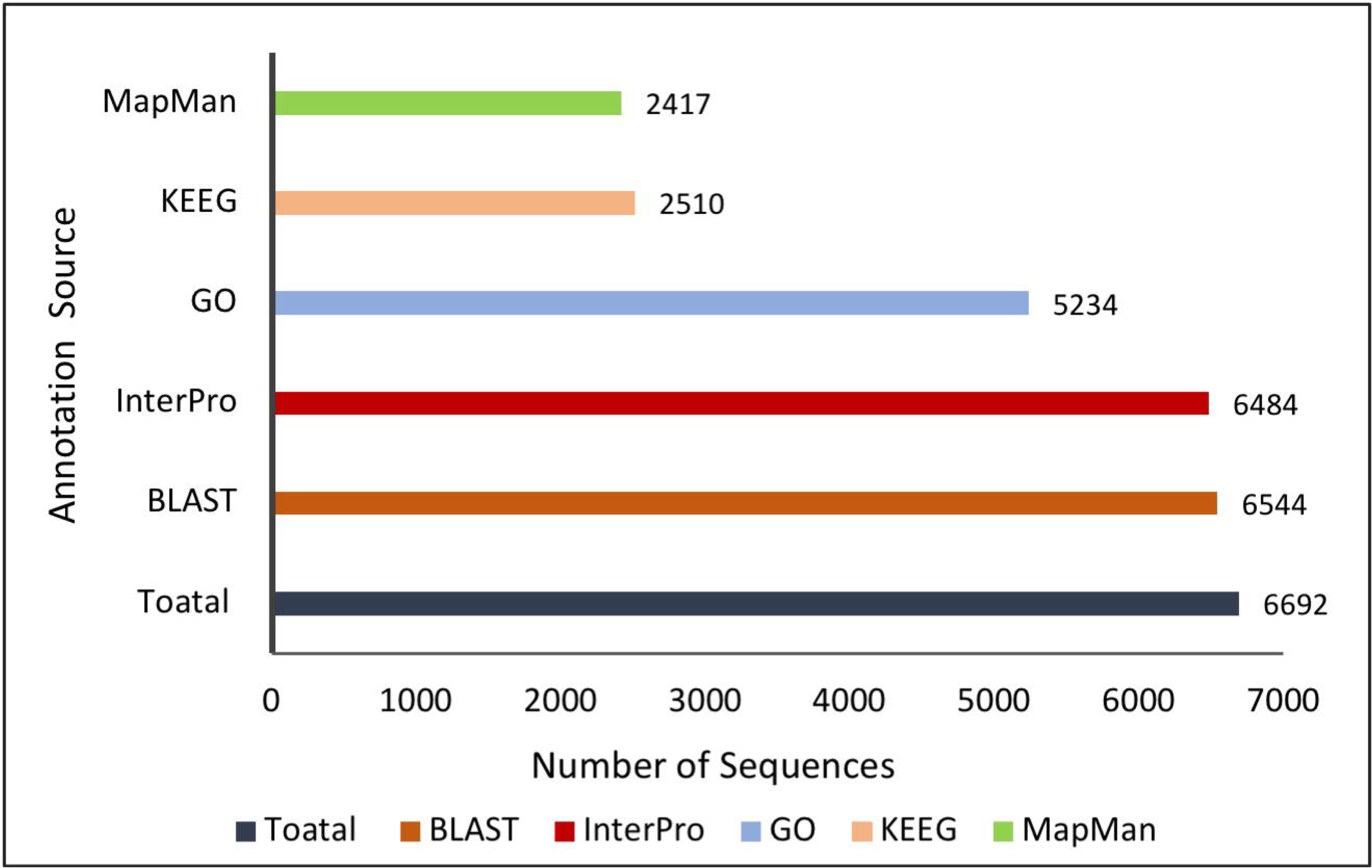
Functional annotation statistics of citrus DETs in response to *P. parasitica* infection, from different databases.

Gene ontology (GO) terms were assigned to 5234 DETs, in which around 41% of GO terms were associated with molecular function (MF), 36% with biological processes (BP), and 22% with cellular components (CC). Within the MF class, most of the DETs were place in the categories of metabolic and cellular processes, response to stimulus and localization. Most represented BP categories were catalytic activity, binding and transport activity (Figure 5). Detailed GO term visualization indicated presence of many cytoskeleton, microtubule, cell wall related terms in all three; BP, MF and CC GO classes (Figure S6). Interactive graphs of most significantly enriched GO terms revealed interlinked cell wall macromolecule and polysaccharide metabolism among biological processes, and oxidoreductase activity and diverse binding related molecular functions (Figure 6). Most of the DETs assigned to oxidoreductase were highly upregulated at both time points (Appendix S2).

**Figure 5.**
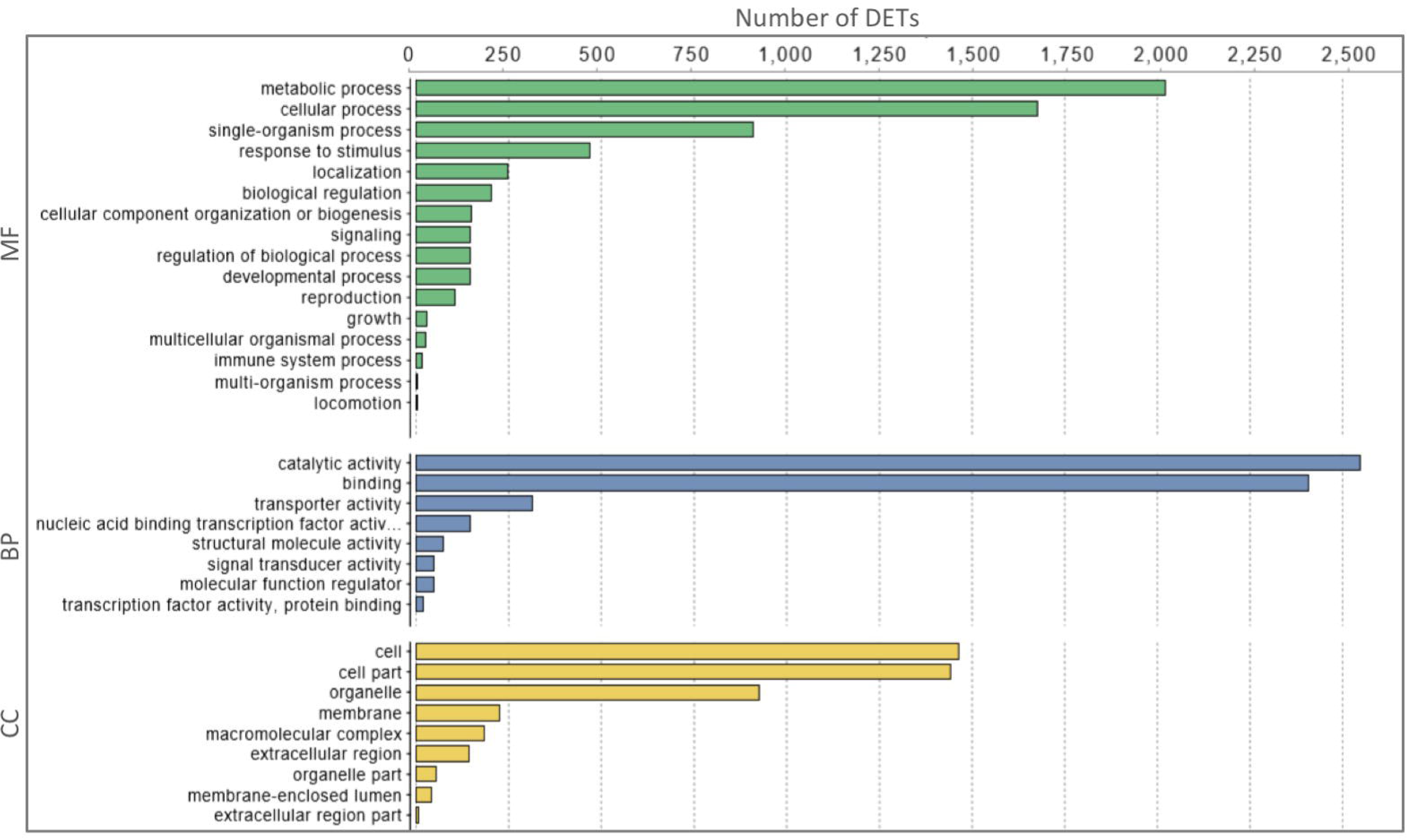
Distribution of GO categories assigned to differentially expressed citrus transcripts in response *Phytophthora parasitica* infection. Number of sequences assigned to top 20 categories in all three classes (molecular function (MF), biological processes (BP) and cellular component (CC)) are shown.

**Figure 6.**
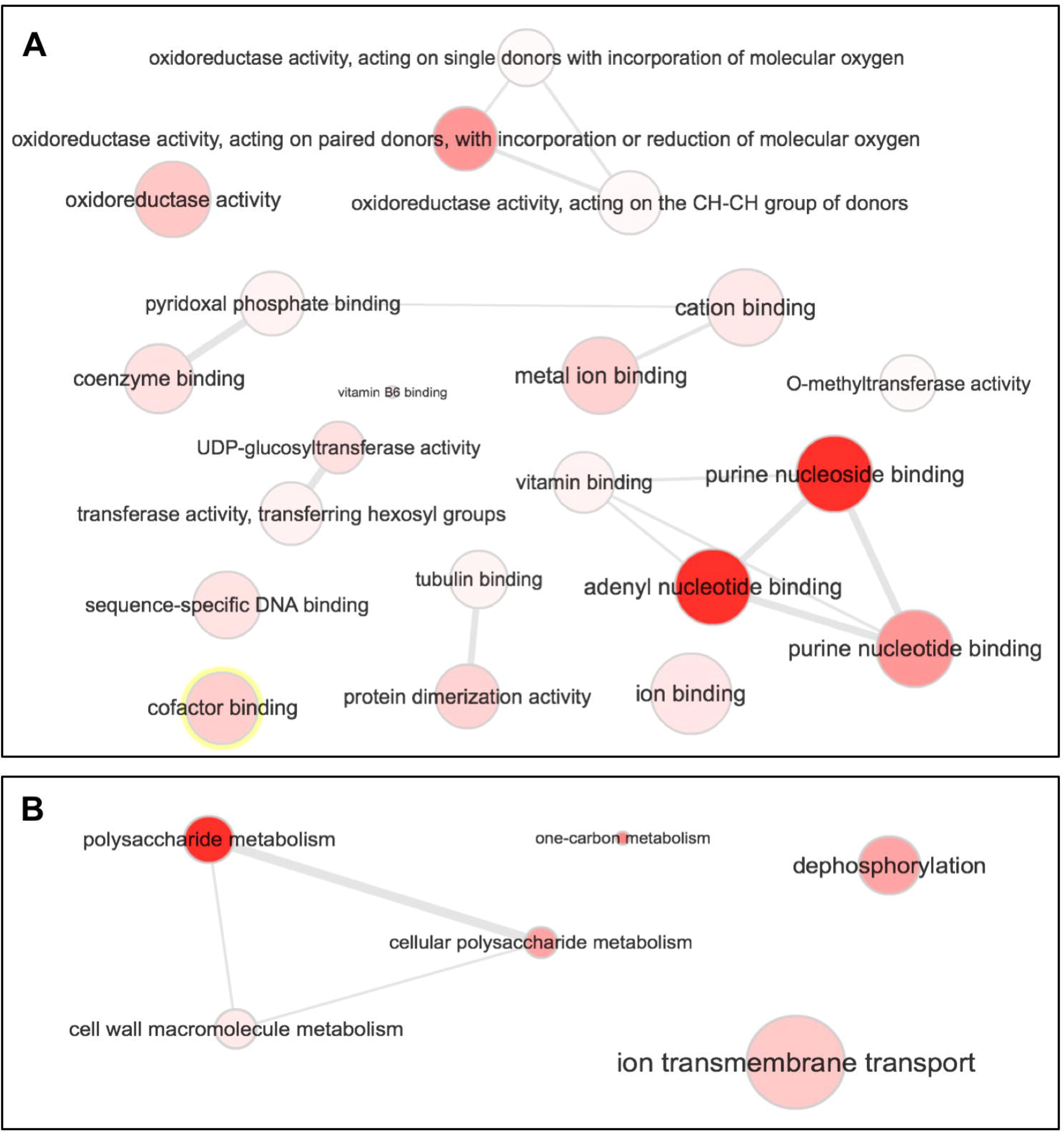
Interactive graphs for summary visualization of most abundant GOs among all significantly enriched GO terms. Most enriched GO terms molecular functions were related to diverse oxidoreductase and binding activities (A). Among biological processes, cell wall macromolecule metabolism, dephosphorylation and ion transmembrane transport were dominant (B). Significance levels are based on enrichment and FDR values with highly significant terms indicated by red nodes. Edges are biologically meaningful links between significant terms.

Cross comparison of daughter terms across 24 hpi and 48 hpi up- and down-regulated DETs revealed some distinct trends. For example, at both time points upregulated DETs were exclusively enriched in transferring phosphorus-containing groups (GO:0016772), protein binding (GO:0005515) and pollen-pistil interactions (GO:0009875 & GO:0048544), whereas, down regulated DETs were more enriched in hydrolase activity (GO:0004553 and GO:0016798), ion binding (GO:0043167, GO:0046872, GO:0046914 & GO:0043169) and lipid localization (GO:0010876). Number of DETs increased from 24 hpi to 48 hpi and so did the enrichment of almost all terms GO terms (Figure S7).

To visualize DETs in biological pathways, transcript sequences were subjected to KEGG annotations and KEGG orthology (KO) terms were assigned to 2510 DETs (Figure 4). The top KEGG pathways for DETs included metabolic pathway, biosynthesis of secondary metabolites, phenylpropanoid biosynthesis and biosynthesis of antibiotics. Plant defense related pathways like Plant hormone signal transduction, Plant-pathogen interaction and MAPK signaling pathway also showed enough mapping of DETs (Table 2). Most of the transcripts mapped on Plant-Pathogen interaction (Figure S8) and MAPK signaling pathway were upregulated at both time points (Figure S9). In plant hormone signal transduction, auxin, cytokinine and gibberellin pathways were all downregulated whereas, ethylene (ET), abscisic acid (ABA), jasmonic acid (JA) and salicylic acid (SA) were mostly upregulated (Figure S10). KEGG enzyme enrichment analysis showed highest mapping on Flavanoid biosynthesis pathway (Figure S11), pathview mapping of DETs on this pathway showed abundance of downregulated genes (Figure S12).

**Table 2.** Top KEGG Pathways for DETs.

| **KO terms** | **Description** | **Number of hits** |
| --- | --- | --- |
| KO01100 | Metabolic pathway | 323 |
| KO01110 | Biosynthesis of secondary metabolites | 192 |
| KO00940 | Phenylpropanoid biosynthesis | 87 |
| KO01130 | Biosynthesis of antibiotics | 85 |
| KO01230 | Biosynthesis of amino acids | 47 |
| KO01200 | Carbon metabolism | 40 |
| KO00500 | Starch and sucrose metabolism | 29 |
| KO04075 | Plant hormone signal transduction | 27 |
| KO04626 | Plant-pathogen interaction | 25 |
| KO04016 | MAPK signaling pathway - plant | 22 |
| KO00941 | Flavonoid biosynthesis | 18 |

Citrus DETs were also visualized in plant functional pathways using the MapMan pathway analysis tool, which accurately assign hierarchal ontologies and provide visual representation of genes involved in different plant processes (Thimm et al., 2004; Usadel et al., 2009).

MapMan bins were assigned to 2417 DETs out of which 468 and 963 were mapped on biotic stress pathway revealing significant enrichment of DETs among all plant defense response categorize at 24 hpi and 48 hpi respectively (Figure 7). In general, most of the genes in cell wall metabolism, abiotic stress, peroxidation and secondary metabolism were upregulated at both 24 hpi and 48 hpi. *PR*-genes, and MAPK signaling were mostly upregulated. Genes in redox state and general signaling were almost equally distributed between the up- and down-regulated categories. Proteolysis was seen more upregulated at 48 hpi compared to 24 hpi. Most of the transcription factors (TFs) like, ERF, bZIP and DOF showed mixed trend of up- and down-regulation whereas most of the WRKY TFs were up-regulated. A notable exception was genes in the MYB transcription family, a majority of which were down-regulated. Consistent to KEEG pathway mappings, plant hormones involved in growth and general physiological processes like auxin and brassinosteroid showed downregulation, and ET and ABA that are generally considered as stress responsive hormones were mostly upregulated (Figure 7 and Figure S10). Overall, these analyses suggested that many different biological, molecular and cellular processes in host are affected by *P. parasitica* infection.

**Figure 7.**
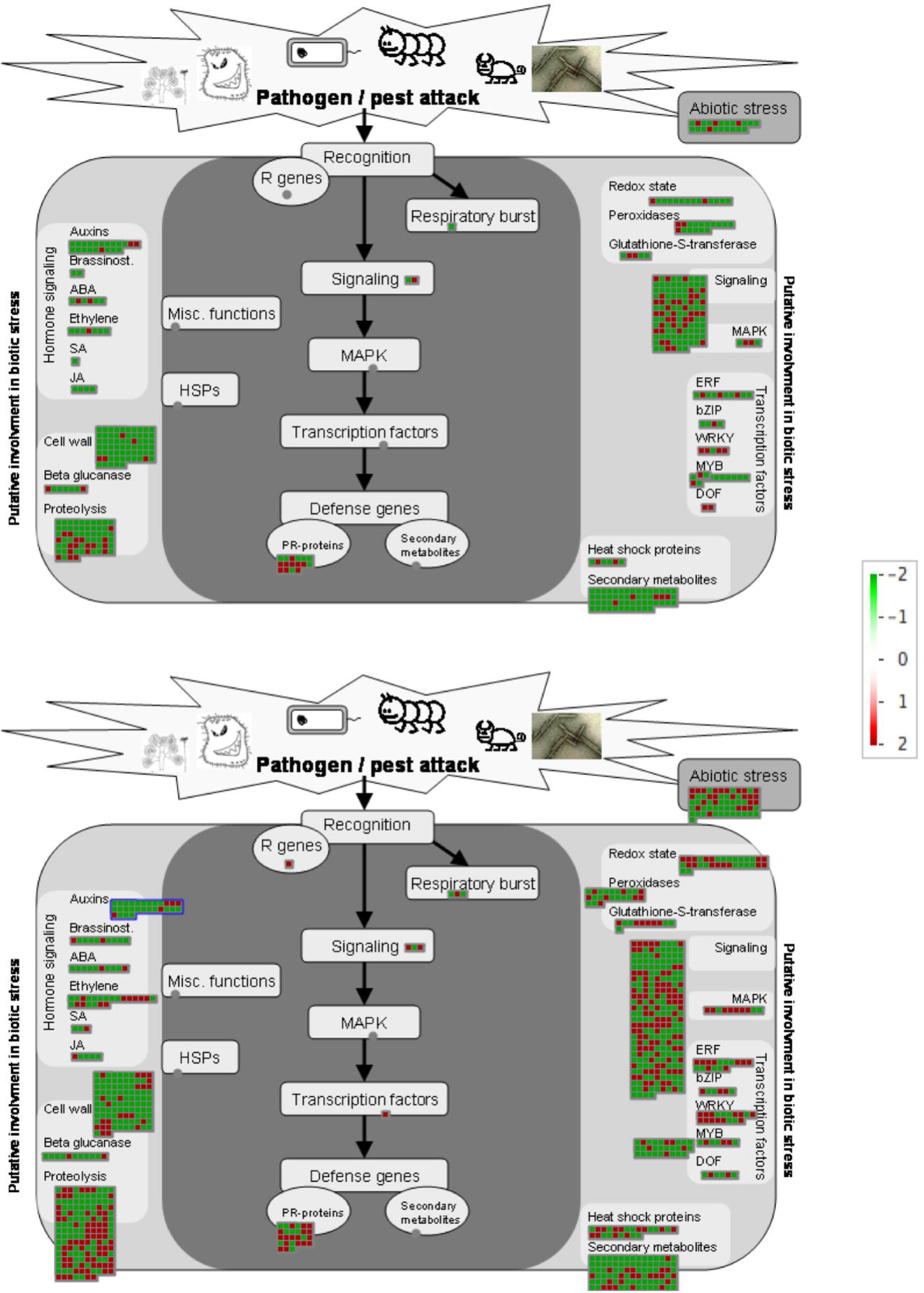
Visualization of differentially expressed genes involved in biotic stress pathway in response to *P. parasitica*. Color gradient represents log2 fold ratios with red representing upregulation and green representing downregulation in treatments over mock roots. Each box represents one transcript. Significant dominance of downregulated transcripts in cell wall, peroxidases, MAPK, heat shock proteins and secondary metabolites are evident. In hormone signaling Auxins, Brassinosteroid and ABA were more downregulated whereas Ethylene and JA categorize are more encompassed by upregulated DETs. In transcription factor categorize WARKY and EFR have more upregulated DETs whereas MYB has more downregulated DETs. Most DETs designated as pathogenesis-related genes (PR-genes) can be seen upregulated. Mixed trends of up and down regulation were seen other biotic stress related processes.

### Putative *R-*genes differentially expressed in citrus roots in response to *P.parasitica*

Functional annotations indicated presence of several *R*-genes among citrus DETs. In addition to that we ran a local BLAST of all citrus transcripts against protein sequences of all *R* genes in Plant resistance genes database (PRGdb). PRGdp is a comprehensive database comprised about 104459 *R* genes from 233 plant species (Sanseverino et al., 2012). In total, 645 transcripts corresponding to 454 genes showed 80-100% (e-value threshold of 1e^-10^) homologies to 490 unique *R* genes. Out of those 490 *R* genes, 284 had proper annotations and 277 of them matched to either *C. sinensis* or *C. clementia R* genes. These putative *R* genes represented all *R* gene classes with 33 containing NB-ARC and LRR domains, which are grouped broadly in NL class in PRGdb, 56 in the CNL class, 12 in the CN, 6 in the TNL, 135 in the RLP, 9 in the RLK (8 RLK-GNK2), 4 in the MLO-like, 27 in the N class and the rest of them were categorized in others class (Appendix S3).

Out of total 454 *R* genes found in the root transcriptome of citrus, 186 were found differentially expressed in response to *P. parasitica* infection. Of those, 100 were upregulated in the range of 4 to 32,768 times and 80 were downregulated (4 to 256 fold change). The top differentially expressed *R* gene was (PRGDB00149304/TRINITY_DN23214_c3_g3) that is a G-type lectin S-receptor-like serine threonine-kinase (SD2-5) was around 8000 times more expressed in response to *P. parasitica* at 24 hpi and reached up to 32000 times upregulated at 48 hpi. Five other *R* genes of G-type lectin S-receptor-like serine threonine-kinases were also found to be upregulated at 1 or both time points. A receptor 12 like uncharacterized *R* gene was more than 2000 times more expressed upon *P. parasitica* infection at both time points. Out of 4 MLO-like genes, one (PRGDB00150044) was upregulated at both time points. Two RGA3 like *R* genes were also found with conflicted differential expression, PRGDB00149184 was downregulated whereas PRGDB00148450 was highly upregulated. Out of 6 TNLs, 3 TMV resistant N-like *R* genes (PRGDB00152395, PRGDB00158343 and PRGDB00149509) were found highly upregulated. Three RPP13 like disease resistance genes were found upregulated; PRGDB00148968 and PRGDB00147909 only at 24hpi whereas PRGDB00149876 was highly expressed at both 24 and 48 hpi. In the CNL class, three different sub types containing of three, four and eight *R* genes that were predictive homologues of At1g12280, At4g27220 and At5g63020 respectively were among the highly upregulated *R* genes (Table 3). Upregulation of several different classes of *R* genes could have a significant role in the preventing disease progression in citrus by *P. parasitica*.

**Table 3:**
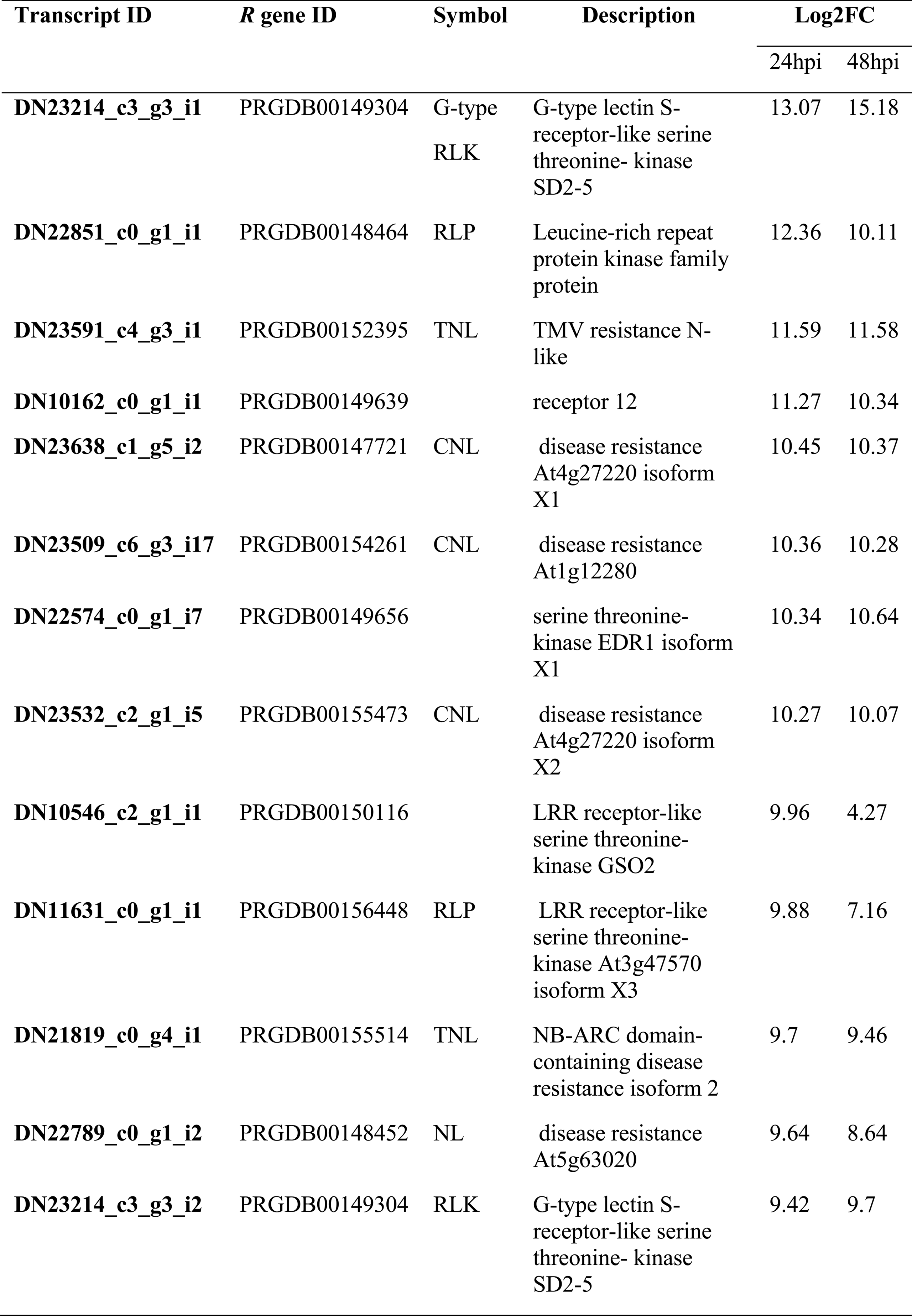
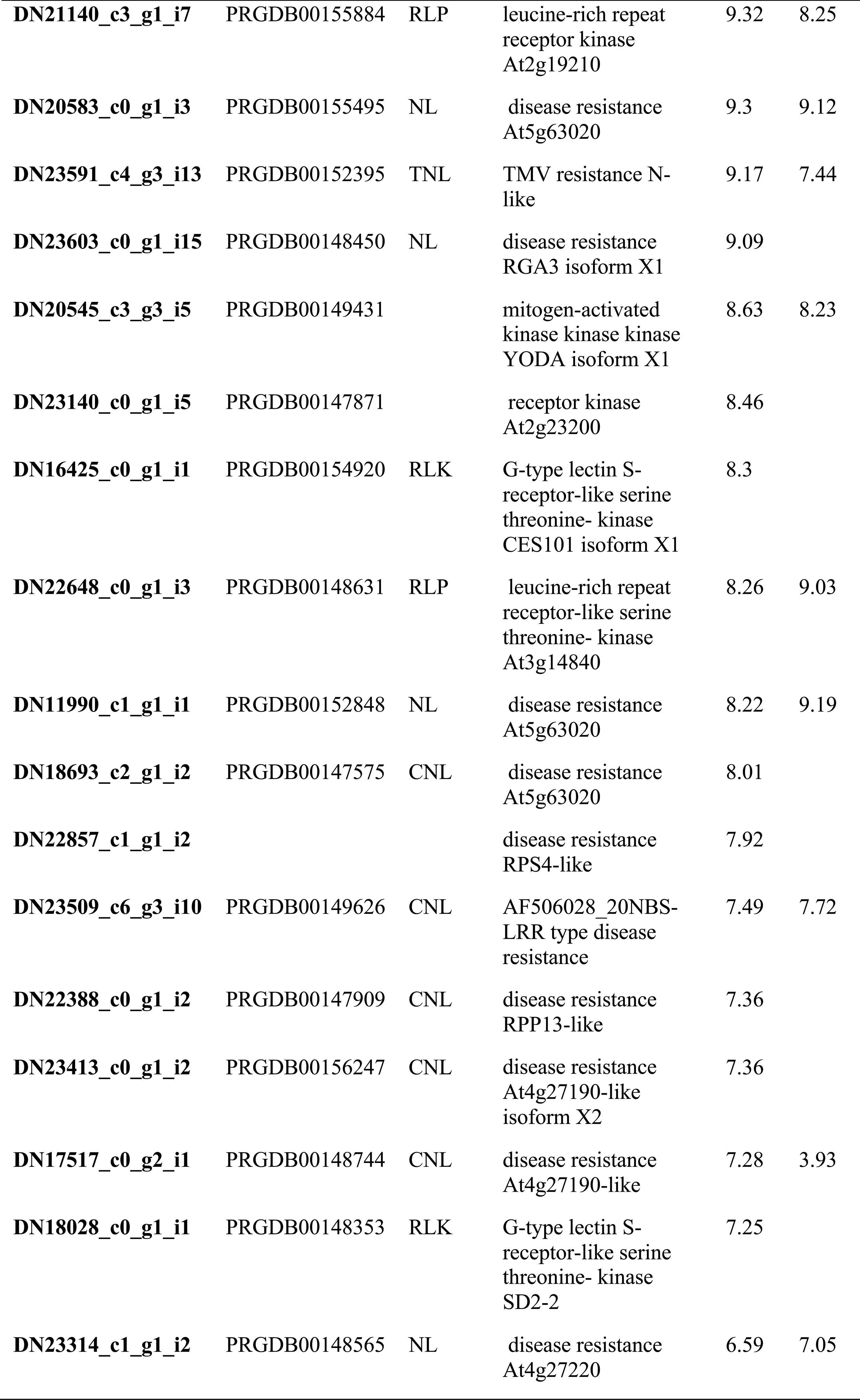
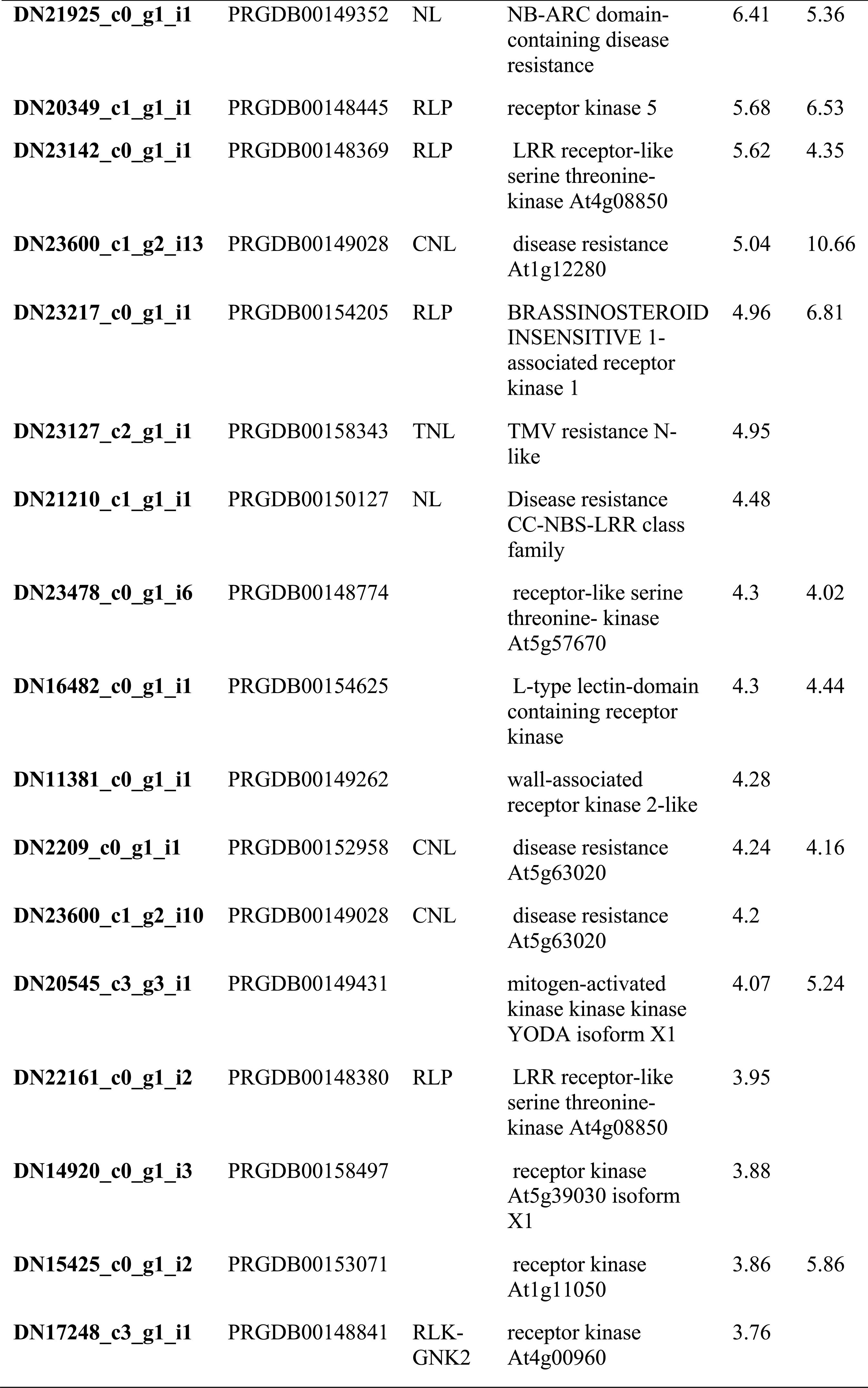
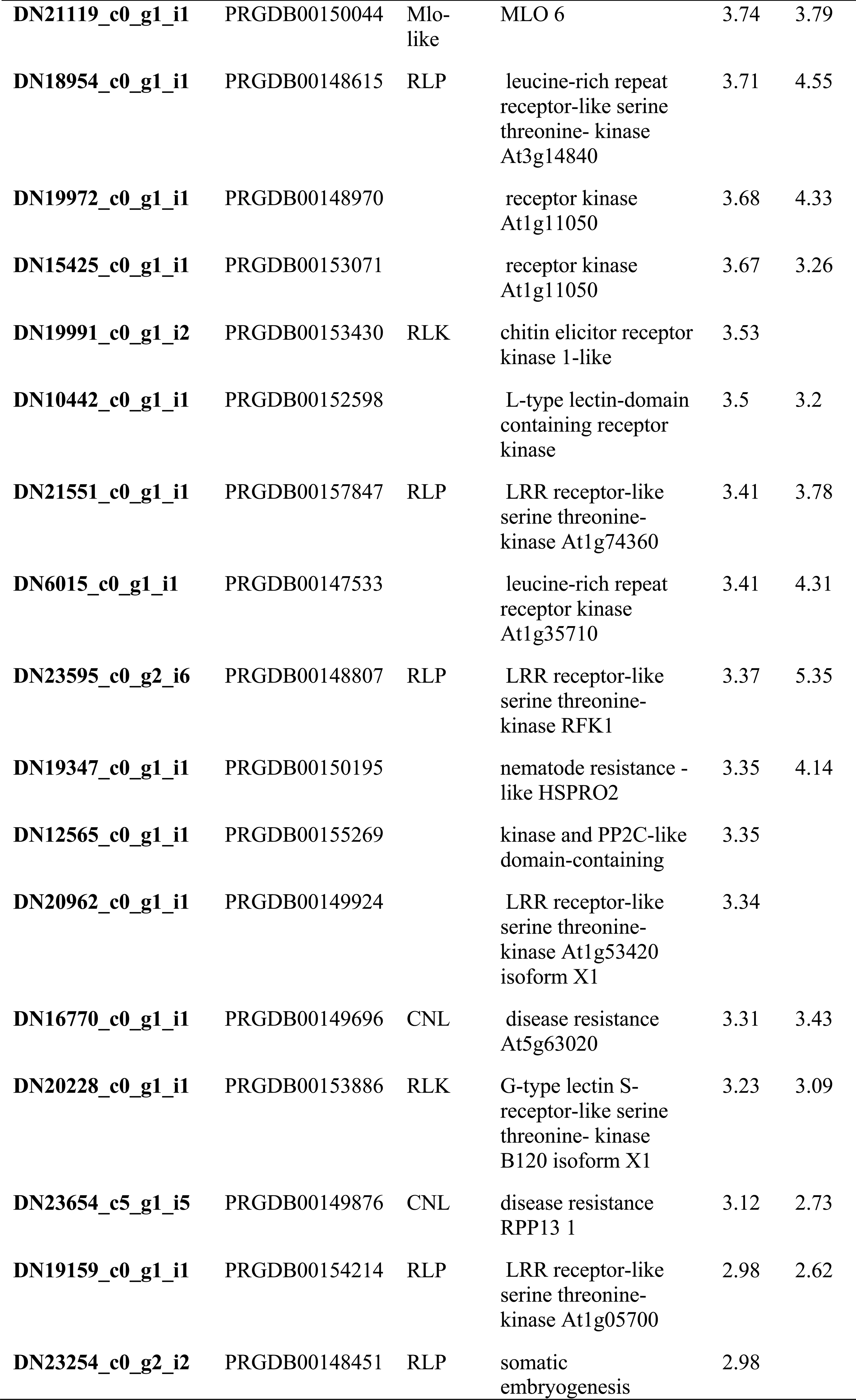
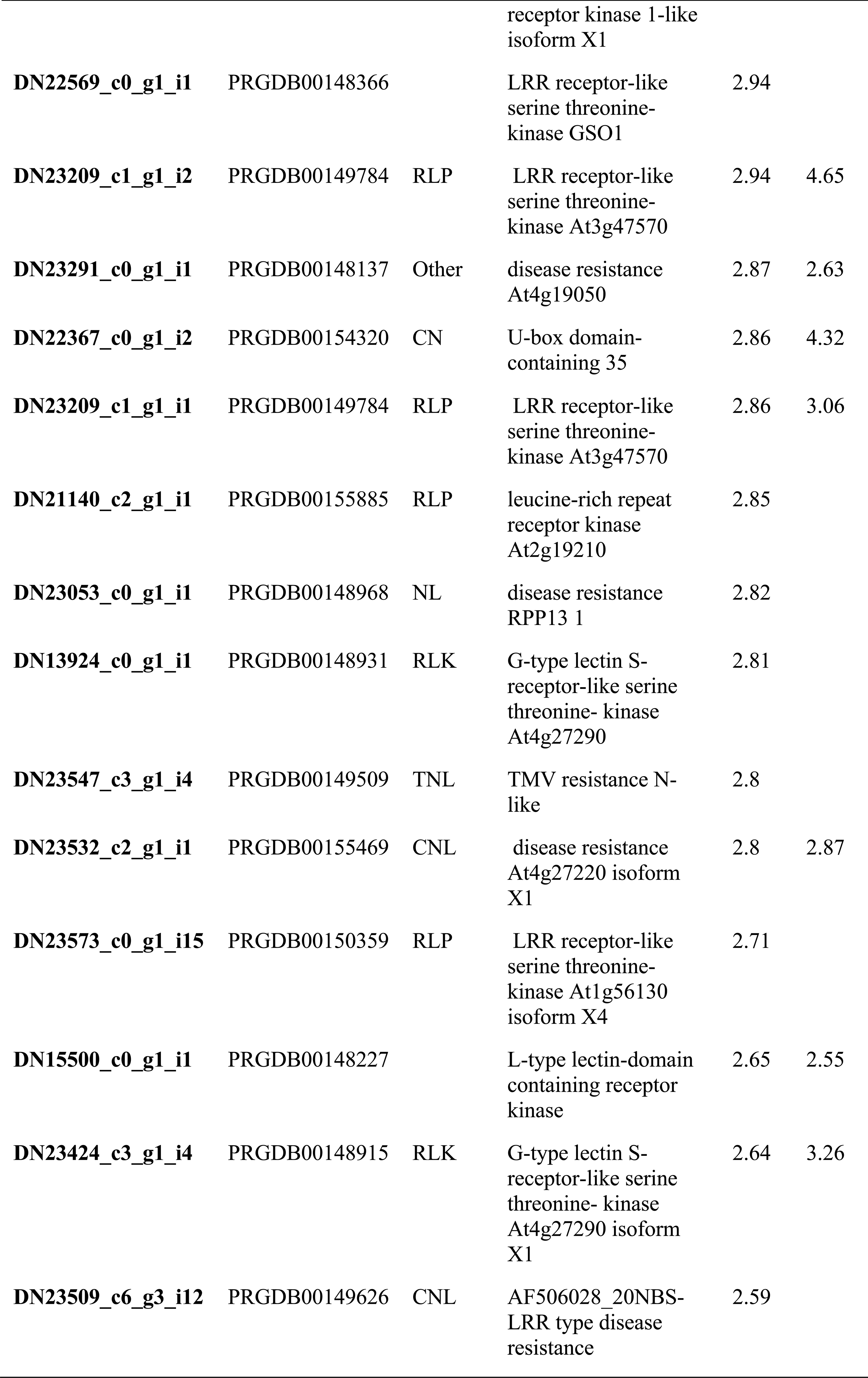
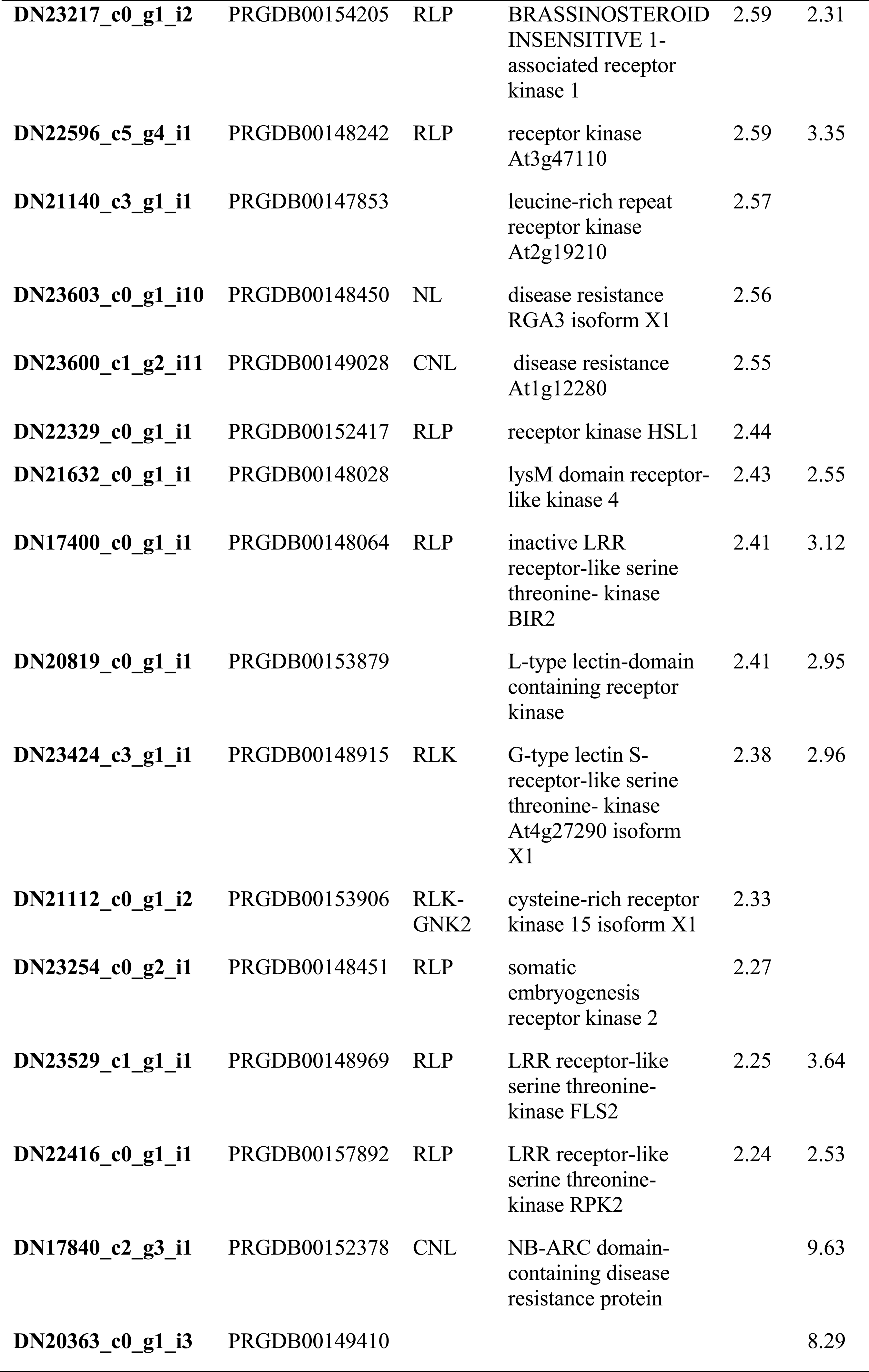
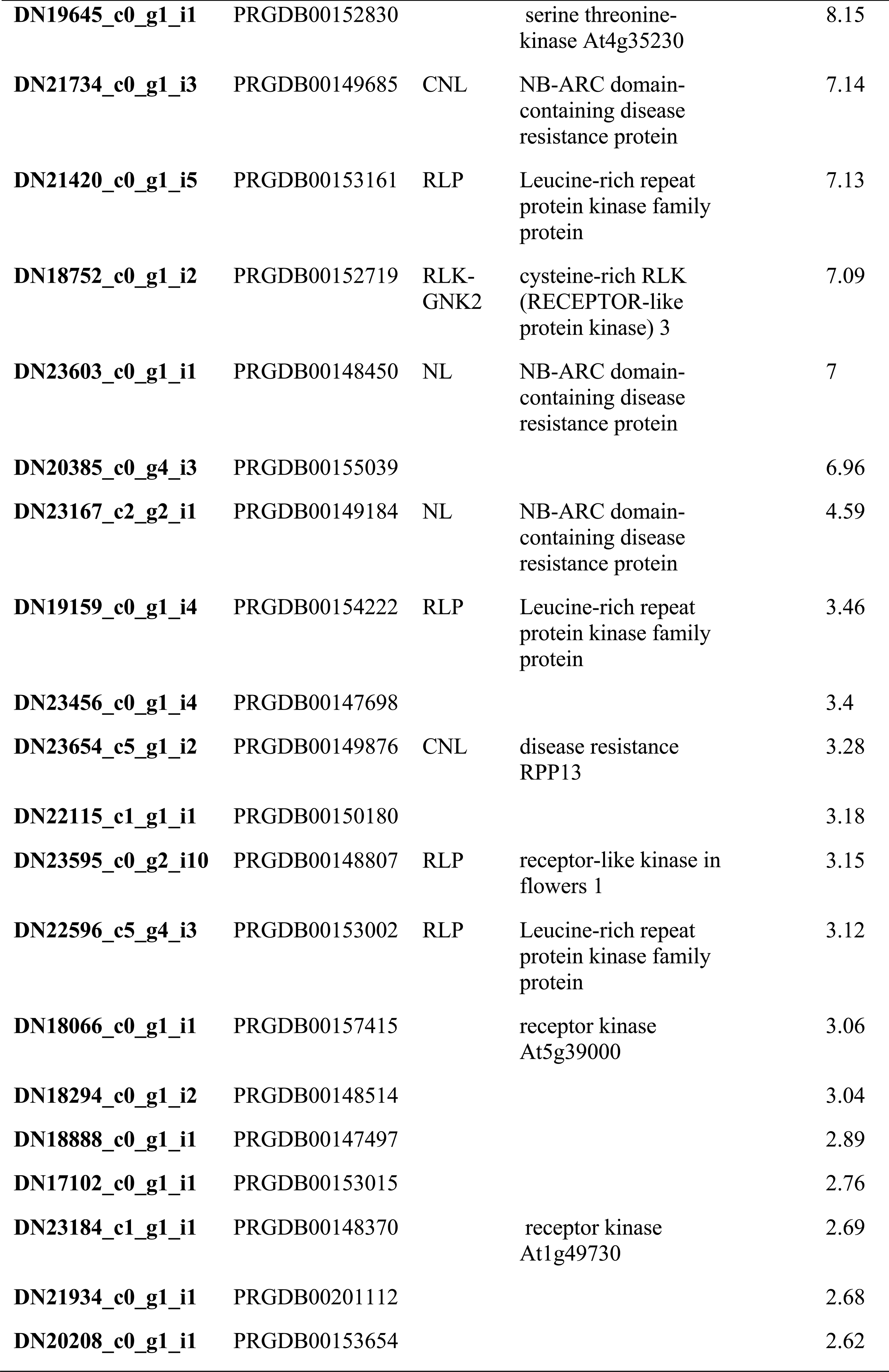
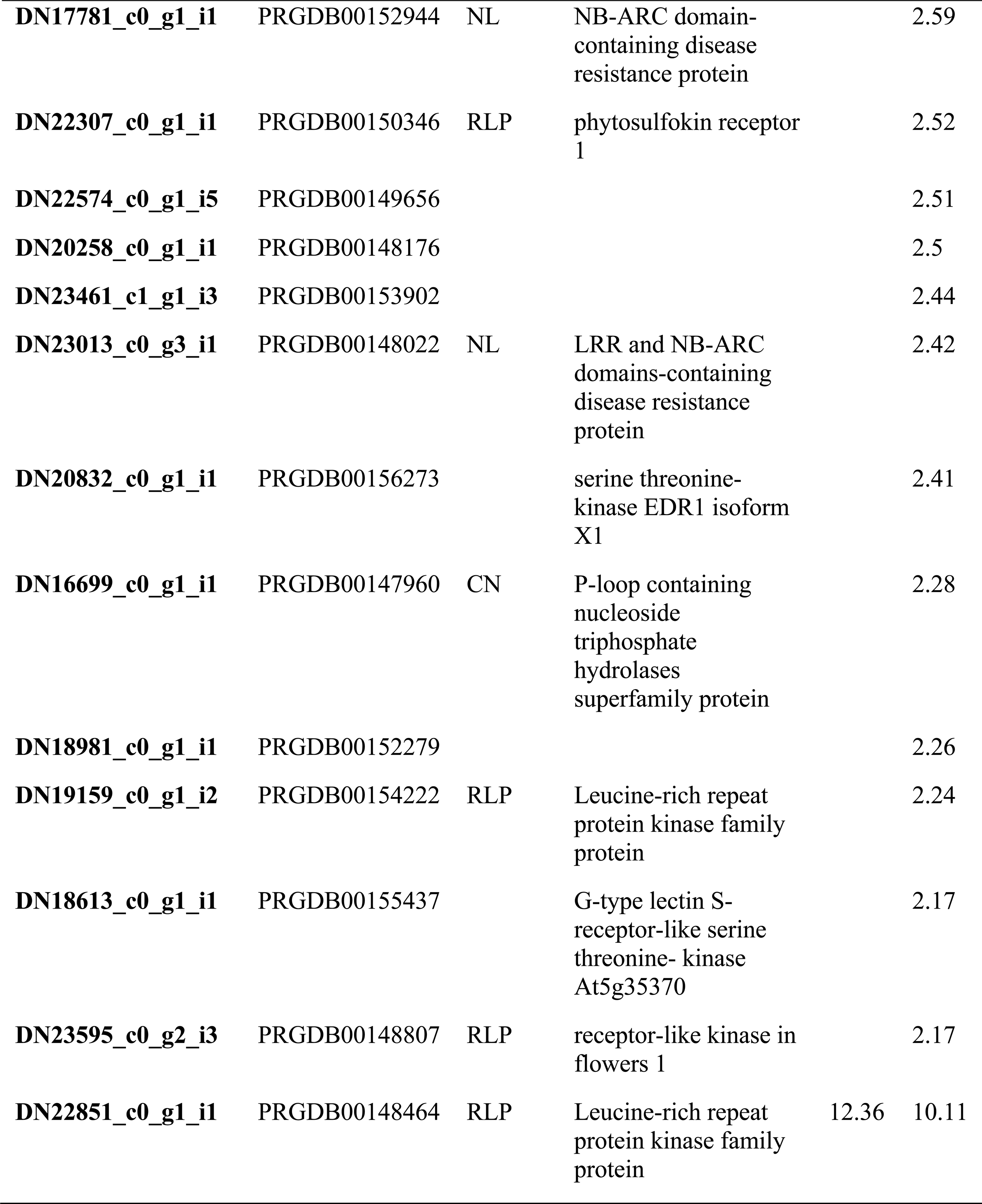
*R* gene DETs induced upon *P. parasitica* infection in citrus roots.

### Validation of RNA-seq results for selected DETs by qRT-PCR

To verify differential expression analysis results from RNA-seq data, transcriptional levels of selected DETs including WRKY transcription factor 31(TRINITY_DN19772_c0_g1_i1), disease resistance At4g27220 isoform X1 (TRINITY_DN23376_c2_g4_i1) and myb family transcription factor APL isoform X2(TRINITY_DN16689_c0_g1_i1), expansin (TRINITY_DN28627_c0_g1_i1), G-type lectin S-receptor-like serine threonine-kinase SD2-5 (TRINITY_DN23214_c3_g3_i1) etc. that were potentially involved in the development of tolerance response in citrus roots against *P. parasitica* and represent different expression profiles across *P. parasitica* treated and control citrus roots samples, were determined by qRT-PCR analysis. Relative expression values of all the tested genes were calculated by using constitutively expressed citrus GADPH gene. Expression profiles of all tested genes resulted from qRT-PCR were in agreement with our RNA-seq data analysis results (Figure 8).

**Figure 8.**
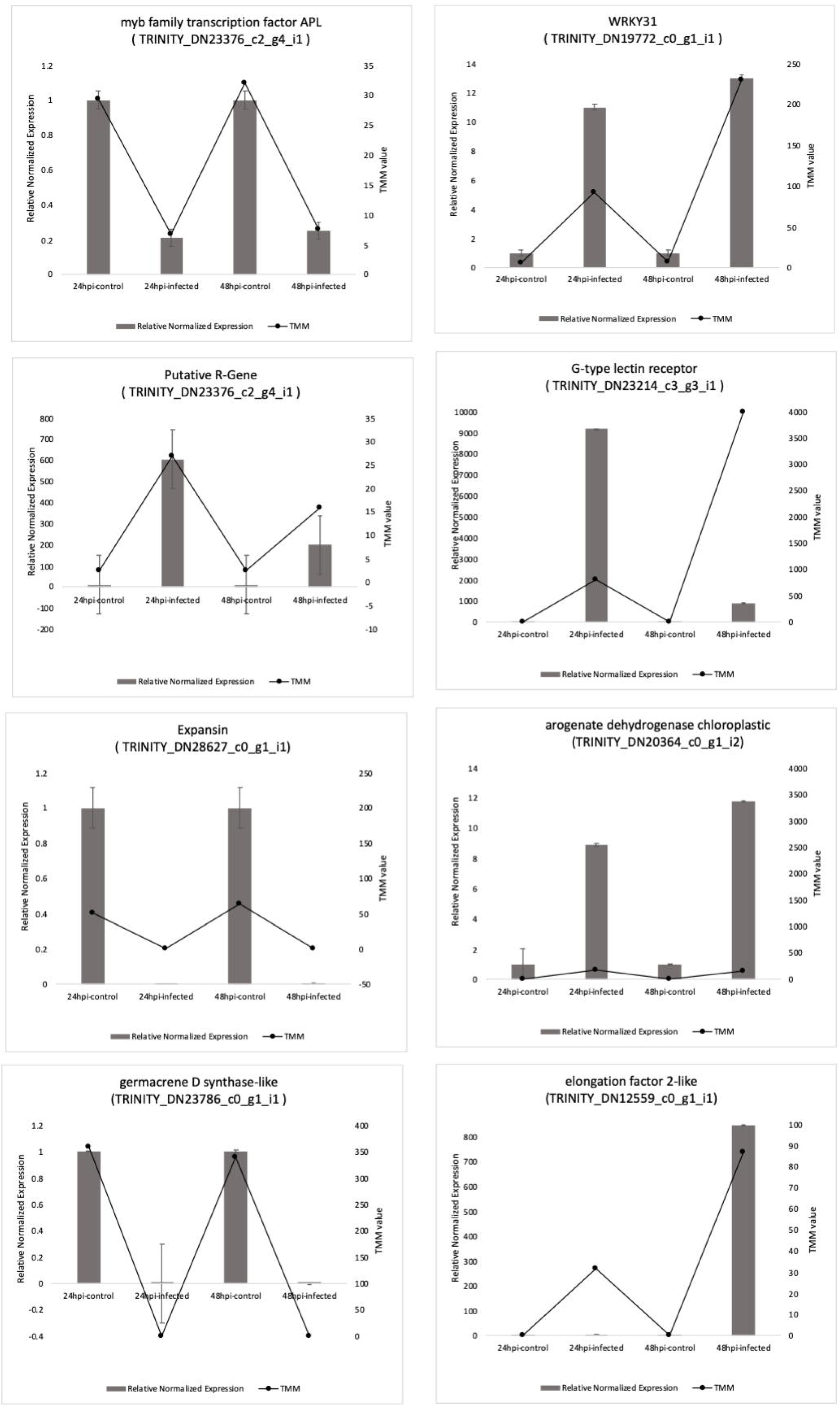
QRT-PCR based validation of plant defense related and top DETs in citrus roots in response to *P. parasitica*. Expression levels of tested genes were normalized based on transcript levels of GADPH gene. TMM values calculated from RNA-seq are compared to relative expression values determined by qRT-PCR analysis. Relative expression values of infected samples were determined against the average values of mock treated samples at each time point. Expression profiles DETs in both analysis correlate well for all the tested genes.

## Discussion

Genetic resistance among wild and cultivated plants against wide variety of phytopathogens are the best resources that can be utilized to develop the most economical and environmentally friendly strategies to fight against *Phytophthora* diseases. In citrus, resistance rootstocks are excellent breeding material for developing *Phytophthora* resistant citrus trees with high quality fruits (Lima et al., 2018). Few rootstocks like Carrizo citrange, Swingle citrumelo and *P. trifoliata* have been reported to be tolerant to *P.parasitica* infection except in disease complexes (Dalio et al., 2017; Graham, 1995; Graham et al., 2003; Hutchison, 1974). Molecular mechanism of resistance of most of these citrus rootstocks to *Phytophthora* pathogens is not well understood. This study was designed to identify *R* genes and overall transcriptional changes in Carrizo citrange rootstock in response to *Phytophthora parasitica* through whole transcriptome sequencing of infected and control samples during early stages of infection.

Microscopic studies of *P. parasitica* treated citrus roots clearly showed the attachment and germination of zoospores on the root surface followed by mycelium colonization (Figure 1), indicating successful initial infection. Mapping coverage of *P. parasitica* genome in the pathogen-inoculated samples increased three times from 24 to 48 hpi (Table S1), which closely corresponded to increased root colonization of pathogen from 24 to 48 hpi observed under the microscope (Figure 1) and is a clear evidence of successful infection and colonization of Carrizo citrange roots by *P. parasitica.* However, disease symptoms were not evident in the above ground parts of the plant suggesting that the understudy rootstock is a host of *P. parasitica* and exhibits a tolerant response unlike *P. trifoliata* that showed non-host resistance (Dalio et al., 2017). To understand the molecular basis of this tolerant response on transcriptome level, we did differential isoform expression analysis in citrus roots during initial stages of *P. parasitica* infection.

A *de novo* transcriptome assembly was generated because read mapping coverage on reference genomes of related *Citrus* spp. (Adhikari et al., 2012; Wang et al., 2014). were quite low (Table S1) which is understood because Carrizo citrange is a hybrid. Assembly optimization was done to keep only biologically meaningful transcripts and to save statistical power in the differential expression analysis (McCann et al., 2017; Ono et al., 2015). Initial differential expression was done by using both coding isoforms counts and unigene counts. Considering the fact that Carrizo citrange is a hybrid and shared exons between two parental genomes could cause ambiguities and significant loss of information in case of unigenes, most of the downstream analysis presented here are based on protein coding transcripts. Although statistically exhaustive and complicated, transcript level differential expression analysis has been proved to provide higher resolution and several new algorithms enable individual transcript abundance estimation (Soneson et al., 2015). In our studies results from both methods were almost similar except the fact that gene-level analysis missed some of the differential expressed transcripts, contrastingly transcript-level analysis covered most of the genes predicted to be differentially expressed by gene-level analysis. Differential expression analysis using transcripts count was also validated by qRT-PCR and expression profiles of all tested genes were found similar to RNA-seq results (Figure 8).

Numerous plant defense related genes putatively involved in preventing diseases progression were identified to be differentially expressed in citrus roots in response to *P. parasitica* infection. In addition to plant defense, major transcriptional reprogramming of diverse cellular and metabolic processes in citrus root upon *P. parasitica* infection was observed. The mechanism of plant’s response to pathogens and development of resistance or tolerant response is a complex phenomenon that involves interconnected network of changes in several regular physiological processes (Jones and Dangl, 2006) that is clearly reflected in transcriptome-based studies of several different plant-pathogen systems(Adhikari et al., 2012; Chen et al., 2014; Gao et al., 2013; Judelson et al., 2008; Kunjeti et al., 2012). Consistent with gene expression profiles of *Nicotiana benthamiana* leaves and *A. thaliana* roots infected with *P. parasitica* during compatible interaction (Le Berre et al., 2017; Shen et al., 2016), transmembrane transport, protein kinase activity, iron ion binding, carbohydrate metabolism, phosphotransferase activity and many other daughter terms of biological and molecular processes were significantly enriched among both up and down regulated DETs in citrus roots. Unlike susceptible *A. thaliana* roots where number of DEGs didn’t increase considerably with time during *P*. *parasitica* infection (Le Berre et al., 2017), total number of DETs increased in citrus roots from 24 hpi to 48 hpi suggesting induction of stronger defense response towards *P. parasitica* overtime that can be related to tolerance response during later stages (Figure 7 and Figure S4).

Upon perception of pathogen generation of reactive oxygen species (ROS) has been reported as one of the earliest defense strategies in plants (Elmayan and Simon-Plas, 2007). Oxidoreductase activity was found to be the most significantly differentially regulated MF in roots upon *P. parasitica* infection and several different oxidases and reductases were found to be highly upregulated. Upregulation in oxidation reduction processes has been observed in resistant response of many plants to different pathogens including *Phytophthora* species (Reeksting et al., 2016). Differential proteome analysis of a resistant soybean cultivar in response to *P. sojae* also revealed significant upregulation in oxidoreductase activity related proteins (QIU et al., 2009). A tobacco NADPH oxidase was found responsible for triggering HR in response to a *Phytophthora cryptogea* elicitor leading to acquired resistance (Elmayan and Simon-Plas, 2007). An NADPH oxidase (TRINITY_DN6244_c1/c0) was 512 times more expressed in citrus roots infected with *P. parasitica* compared to control (Appendix S2) so it might be involved in ROS signaling during early stages of infection but that was not enough to prevent pathogen colonization. *P. parasitica* is a hemi biotroph so is adapted to survive oxidative burst during initial stages of infection (Attard et al., 2010).

After perception and attachment to the host’s surface, the next step for successful infection is penetration into the host. Cytoskeleton reorganization of infected cells is considered an important component of plant defense against pathogen penetration (Hardham et al., 2007; Underwood, 2012; Higaki et al., 2011). Significant enrichment of membrane and cell wall macromolecule metabolism among DETs (Figure 6), and representation of microtubule and overall cytoskeleton organization among all three GO (MF, BP & CC) classes were observed (Figure S6). In addition to that enrichment of cell wall related genes in the biotic stress overview of MapMan is also evident (Figure 7). Expansins are reported to play an important role in plant growth and involved in cell wall loosening, disassembly and separation during cell growth. We found differential modulation of 29 different expansins at both extremes of up and downregulation. For instance, expansin B 1 (TRINITY_DN21518_c1_g2_i8) was 3565 time more expressed and unclassified expansin (TRINITY_DN28627_c0_g1_i1) was 304 times downregulated in *P. parasitica* infected roots as compared to uninfected roots (Appendix S2). Unlike our results only downregulation of four expansins was seen during the compatible interaction of *P.parasitica* in *A. thaliana* roots (Le Berre et al., 2017). Most of the other cell wall and microtubule related genes were significantly down regulated in the infected cells, which could be either an attribute of plant defense response against pathogen infection or pathogen’s strategy to manipulate host cell for successful penetration.

Consistent with the findings of Le Berre *et al*. in *A. thaliana* roots, some VQ motif contacting genes were found to be upregulated in citrus roots during early stages of *P. parasitica* infection (Le Berre et al., 2017) but at 48 hpi mix trend of up and down regulation was seen (Appendix S2. Unlike their finding, VQ29 reported to be specifically involved in resistance against *P. parasitica* was not found differentially expressed in our study (Le Berre et al., 2017).

Phytohormones, specifically SA, JA and ET are considered an integral part of immune response (Denance et al., 2013), which lead to broad spectrum systemic acquired resistance (SAR) and induced systemic resistance (ISR) (Shoresh et al., 2010). SA is considered an important component of SAR, which involves induction of numerous *PR* genes not only in the infected cells but also in non-infected systemic tissues (Durrant and Dong, 2004). Another important component of SAR, downstream to SA is NPR1 (Vanacker et al., 2001), which interacts with TGA transcription factor of the bZIP family to induce expression of *PR* genes ultimately giving rise to SAR (Durrant and Dong, 2004). Some *PR* genes, NPR homologs, one NPR suppressor and three TGA transcription factors were differentially modulated in citrus roots in response to *P. parasitica* infection. Both JA and ET are more closely associated with ISR, usually antagonistic to the SA pathway, and the balance between these three signaling pathways is determined by the lifestyle of the pathogen (Dong, 1998; Heil and Bostock, 2002). DETs involved in ET pathway were mostly upregulated in our studies, whereas, mixed trend of up and down regulation was observed in the putative JA DETs (Figure 9). Foliar application of JA in potato and tomato have been shown to induce resistance against *Phytophthora infestans* (Cohen et al., 1993). In case of *N. benthamiana-P. parasitica* interactions, induction of JA and ET pathway is reported but SA pathway was not seen to be activated (Shen et al., 2016). Plants generally activates JA/ET pathway and repress the SA pathway infected by necrotrophic pathogens. In contrast, activation of SA and repression of JA/ET pathways have been observed in response to biotrophic pathogens (Spoel et al., 2007). In contrast to our results, hormonal cross talk was shown not to be involved in resistant response of *P. trifoliata* to *P. parasitica* (Dalio et al., 2017). Flavonoid biosynthesis that is also linked to plant hormones regulation especially SA pathway, was also found highly enriched differentially modulated genes (Figure S11 and S12). Differential induction or repression of JA, ET and SA pathways in citrus roots in response to *P.parasitica* infection represents a cross talk of these defense hormones in dealing with the hemi-biotrophic nature of *P. parasitica* infection, which might be responsible for the host’s tolerance response to disease despite heavy pathogen colonization of roots.

Functional annotations revealed DETs involved in reproduction (Figure S6) seven of them were found with pollen allergen signatures (Appendix S2). These pollen-pistil interaction proteins are generally known as allergens as their homologs in many plants cause allergies in animals (Rogers et al., 1991; Rogers et al., 1992). They are characterized as pectate lyase enzymes that might have some role in cell death of unwanted reproductive structures during fruit development (Wu and Cheung, 2000). Role of pectate lyase in deterioration of host cells during infection has also been reported in host-pathogen interaction studies but most of the known pectate lyases involve in infection are of pathogen origin (Vorwerk et al., 2004). Recently, Mondragon et al (2017) have reported some transcriptional level resemblances in *Arabidopsis* flower in response to fungal infection and pollination. They have observed similar transcriptional trends in some genes putatively involved in pollination and immune response to *Fusarium graminearum* that penetrates through pistils of *Arabidopsis* flowers (Mondragon et al., 2017). Differential expression of pectate lyase genes in roots may have some role in HR or pollen pistil interactions might have some association between plant response to intracellular hyphal penetration and pollen tube penetration phenomenon in flowers.

*R* genes are an integral component of host-pathogen interactions, involved in sensing pathogen patterns (RLPs and RLKs) and pathogen-induced manipulations in host cells followed by triggering an array of responses including local cell death or hypersensitive response (HR), phytohormone signaling, activation of *PR* genes and many other defense related pathways (Thomma et al., 2011; Wanderley-Nogueira et al., 2012). More than 40% of the total *R* genes identified in citrus root transcriptome were found to be differentially modulated upon *P. parasitica* infection. Significant induction of several important *R* gene classes was observed that could be putatively involved in tolerant response to *P.parasitica* (Table 3 and Appendix S3).

Infection of *P. parasitica* in *N. benthamiana* leaves caused up-regulation of 56 LRR-RLKs (Shen et al., 2016). We found upregulation of around 68 RLPs and RLKs indicating significant induction of pathogen sensors in citrus roots in response to *P.parasitica* infection. Lectin receptor kinases are an important class of RLKs, comprising of three (G-type, L-type and C-type) further subclasses. Six G-type RLKs including the top induced *R* gene (32000 times upregulated) were induced in citrus roots upon *P. parasitica* infection compared to control (Table 3). L-type RLKs contribute in plant defense and reported to confer resistance against *Phytophthora* species (Wang and Bouwmeester, 2017; Wang et al., 2015) but G-type has never been reported as *Phytophthora* resistant gene. In rice *BpH3*, a resistance gene comprises of three G-type RLKs is known to provide broad spectrum stable resistance against rice brown plant hopper since last thirty years. *Pi-d2* another G-type RLK in rice confer resistance to rice blast fungus *Magnaporthe* grisea (Wang et al., 2018). Receptor 12 was another uncommon uncharacterized *R* gene that was highly responsive to *P. parasitica* in citrus roots (Table 3). Whether, these receptors play a role in resistance to *Phytophthora* would be an interesting to investigate in future studies.

Boava *et al*. (2011) have reported upregulation of two *R* genes (one TNL and one RPS4 type) in *P. parasitica* resistant *Poncirus trifoliata* plants compared to *Citrus sunki* genotypes, which are reported to be susceptible to *P. parasitica* (Boava et al., 2011). Consistent to their findings, one RPS4 homologue (TRINITY_DN22857_c1_g1) and four TNLs including three TMV resistant N-like genes were among the highly induced *R* genes in citrus against *P. parasitica*. In a late blight resistance potato cultivar SD20, TMV and RGA3 type resistant genes were found upregulated in response to *P. infestans* (Yang et al., 2018). One RGA3 homologue was induced and one depressed in our studies. RPP13 locus of *A. thaliana* has been reported to offer broad spectrum resistance against different strains of a biotrophic oomycete pathogen *Peronospora parasitica* that cause downy mildew in different plants (Bittner-Eddy et al., 2000). Three RPP13 homologs were found upregulated in citrus roots in response to *P. parasitica*. Apart from these examples, several other genes belong to different classes of *R* genes were highly induced pathogen challenged citrus roots. Involvement of these *R* genes along with other defense regulators have been suggested to play a significant role in preventing disease progression in Carrizo citrange in response to *P. parasitica*. Transcriptional reprogramming of most of the cellular and physiological processes and all components of biotic stress responses especially high induction of several *R* genes, modulation of phytohormone signaling, proteolysis, TFs and signaling in Carrizo roots indicates significant activation of defense during infection that increase overtime and thus prevent diseases progression after colonization and confer a tolerant response against *P. parasitica.*

## Acknowledgments

This work was supported by a grant to G.S.A. by the Institute of Food and Agriculture Sciences, University of Florida. Z.A.N. was supported by a grant by the Institute of International Education, Fulbright Pakistan.

## Supplementry Figures

**Figure S1. Plot showing ExN50 statistics.** E90N50 value indicates 90% of expressed transcripts have N50 value of 2056bp that indicates a very good quality assembly.

**Figure S2. Pipeline of RNA-Seq data analysis.**

**Figure S3. Volcano plots showing pairwise comparisons between *P. parasitica* infected and mock-treated control roots samples.** Dispersion of total differentially expressed transcripts found in: 48 hpi infected roots Vs. 24 hpi infected roots comparison (A), 24 hpi infected roots Vs 24 hpi mock treated roots transcriptome comparison (B) and 48 hpi infected roots Vs 48 hpi mock treated roots comparison (C). Log fold change values are plotted at X-axis Vs. False Discovery Rate (FDR) at the Y-axis. Red color is indicating transcripts with statistically valid FDR values.

**Figure S4. Venn diagrams shown number of citrus DETs**. Total DETs (*P*-value < 0.001 and log_2_ fold change > 2) across all comparisons are shown in (A) Upregulated DETs are shown in (B) and Downregulated DETs are shown in (C). Pink circle represents 24hpi treatment vs 24hpi mock comparison, green circle is 48hpi treatment vs. 48hpi mock and yellow is 48hpi treatment vs 24hpi treatment.

**Figure S5: Top hit species distribution graph.**

**Figure S6: Detailed GO term annotations of DETs** in biological processes, molecular function and cellular component classes.

**Figure S7. Snap shot of SEACOMPARE.** It is Cross comparison of Singular Enrichment Analysis (SEA).

**Figure S8. Visualization of differentially expressed transcripts involved in MAPK signaling pathway.** Color gradient represents log2 fold ratios with red representing upregulation and green representing downregulation in treatments over mock roots. Left and right-hand side of each KO box show DETs mapping at 24hpi and 48hpi respectively. White box or no color fill means no DET was assigned to that KO term. Half white box on either side means that term was not differentially expressed at the respective time point. Significant dominance of up regulated transcripts was seen in whole pathway.

**Figure S9. Mapping of differentially expressed transcripts on Plant-pathogen interaction pathway.** Color gradient represents log2 fold ratios with red representing upregulation and green representing downregulation in treatments over mock roots. Left and right-hand side of each KO box show DETs mapping at 24hpi and 48hpi respectively. White box or no color fill means no DET was assigned to that KO term. Half white box on either side means that term was not differentially expressed at the respective time point.

**Figure S10. Differentially expressed transcripts mapped on Plant hormone signaling pathway.** Color gradient represents log2 fold ratios with red representing upregulation and green representing downregulation in treatments over mock roots. Left and right-hand side of each KO box show DETs mapping at 24hpi and 48hpi respectively. White box or no color fill means no DET was assigned to that KO term. Half white box on either side means that term was not differentially expressed at the respective time point.

**Figure S11. KEGG enzyme enrichment analysis showed maximum mapping on Flavanoid biosynthesis pathway.** Color key represents names of different enzymes involved in the pathway.

**Figure S12. DETs mapped on Flavanoid biosynthesis pathway.** Color gradient represents log2 fold ratios with red representing upregulation and green representing downregulation in treatments over mock roots. Left and right-hand side of each KO box show DETs mapping at 24hpi and 48hpi respectively. White box or no color fill means no DET was assigned to that KO term. Half white box on either side means that term was not differentially expressed at the respective time point.

## Appendices

**Appendix S1.** Quality report and trim report.

**Appendix S2.** List of transcripts (DETs) that were differentially expressed between pathogen treated roots and mock inoculate roots at 24hpi and 48hpi along with their functional annotation.

**Appendix S3.** Transcripts annotated as *R* genes.

