## Supplementary Figures for "Transcriptome profile of Carrizo citrange roots in response to *Phytophthora parasitica* infection"

e-Xtra

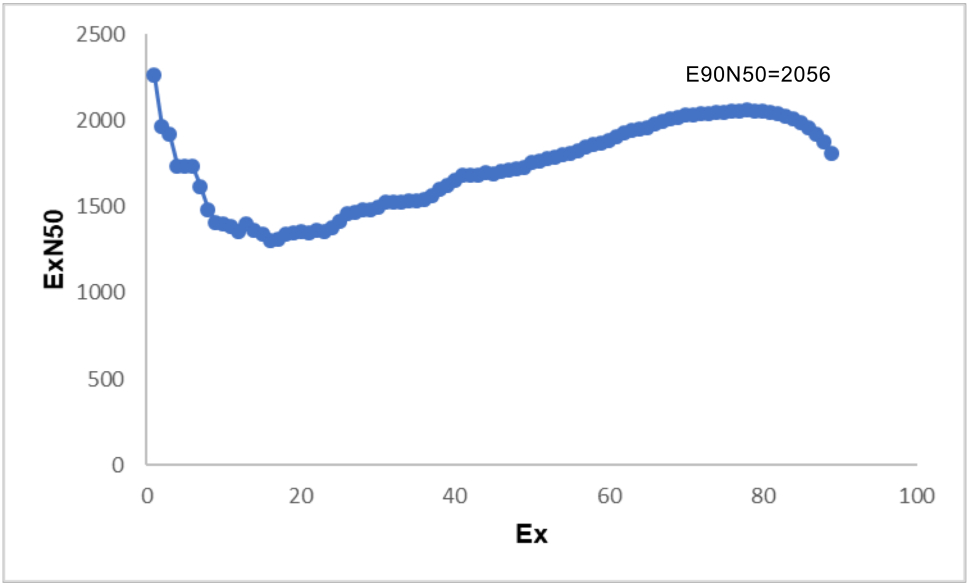


**Figure S1. Plot showing ExN50 statistics.** E90N50 value indicates 90% of top highly expressed transcripts have N50 value of 2056bp that indicates a very good quality assembly.

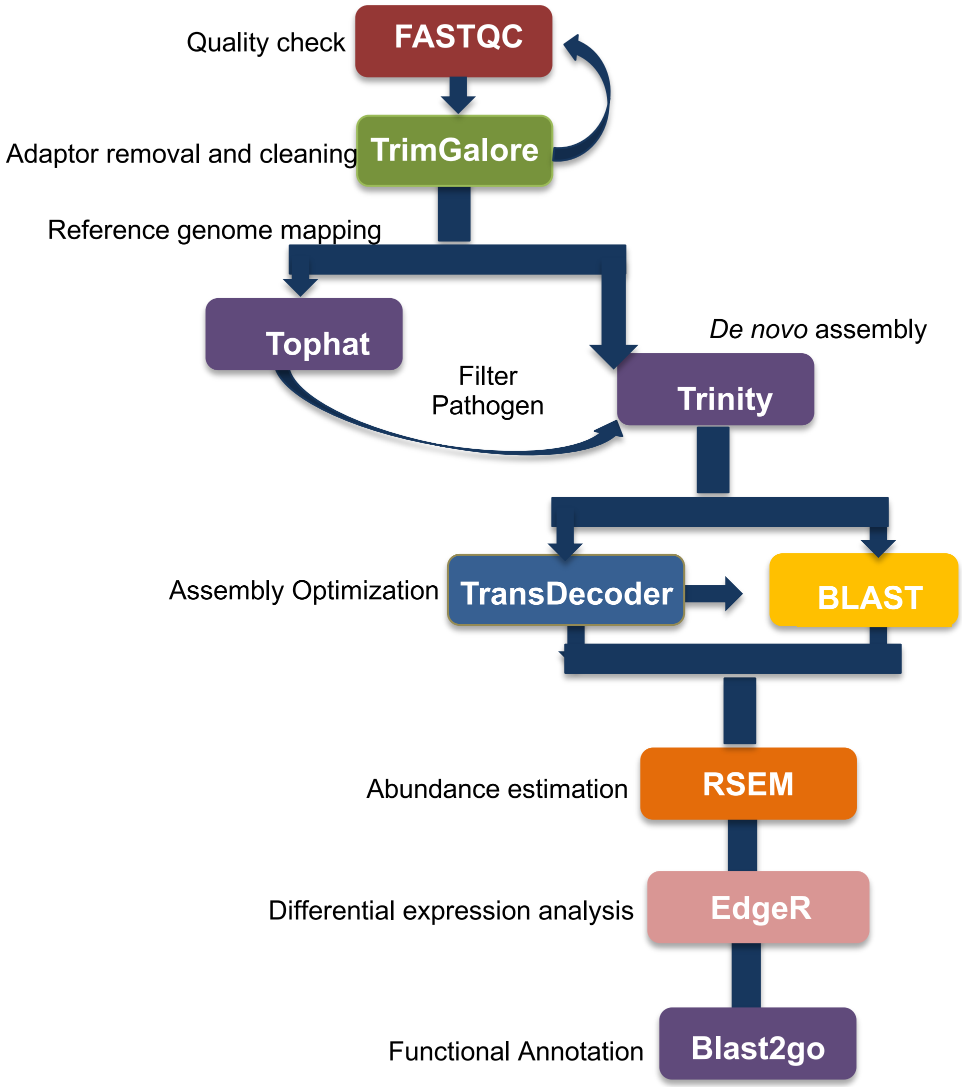


**Figure S2.** Pipeline of RNA-Seq data analysis.

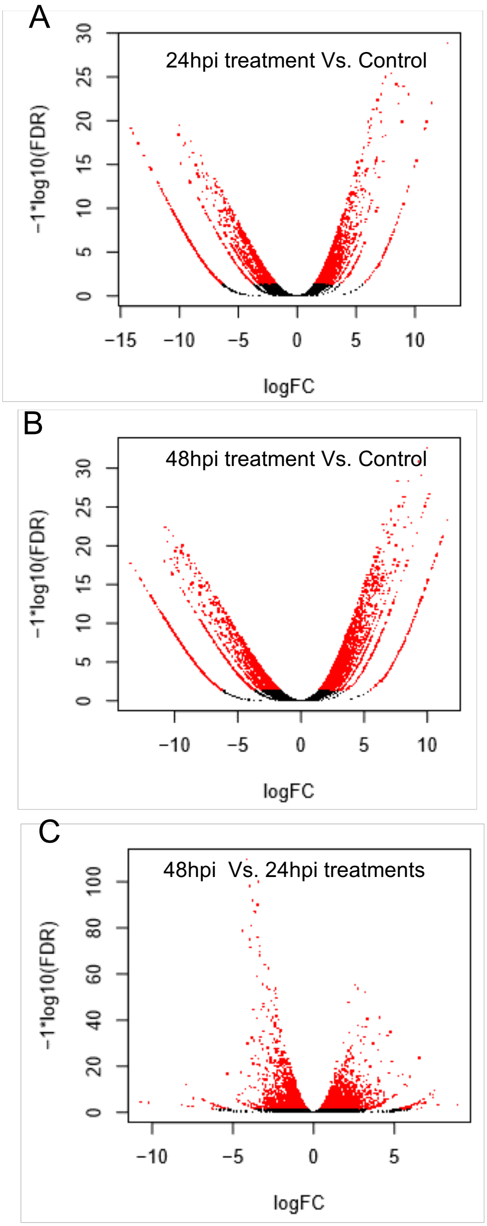
A
24hpi treatment Vs. Control
−1*log10(FDR)
0
5
10
15
20
25
30
−15
−10
−5
0
5
10
logFC
B
48hpi treatment Vs. Control
−1*log10(FDR)
0
5
10
15
20
25
30
−10
−5
0
5
10
logFC
C
48hpi Vs. 24hpi treatments
−1*log10(FDR)
0
20
40
60
80
100
−10
−5
0
5
logFC

**Figure S3. Volcano plots showing pairwise comparisons between *P. parasitica* infected and mock-treated control roots samples.** Dispersion of total differentially expressed transcripts found in: 48 hpi infected roots Vs. 24 hpi infected roots comparison (A), 24 hpi infected roots Vs 24 hpi mock treated roots transcriptome comparison (B) and 48 hpi infected roots Vs 48 hpi mock treated roots comparison (C). Log fold change values are plotted at X-axis Vs. False Discovery Rate (FDR) at the Y-axis. Red color is indicating transcripts with statistically valid FDR values.

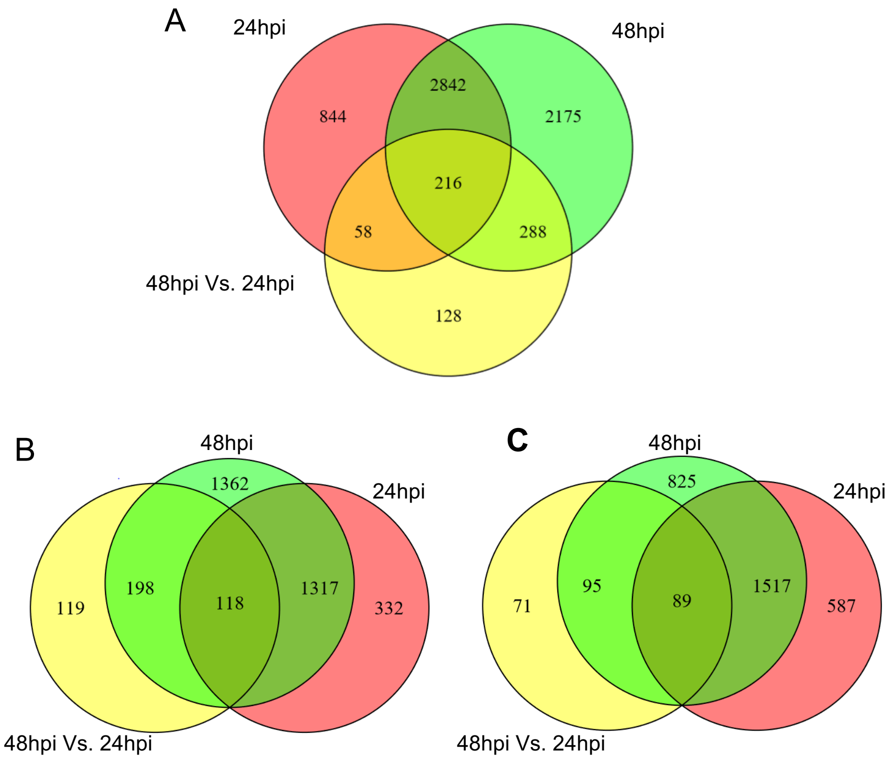


**Figure S4. Venn diagrams shown number of citrus DETs**. Total DETs (*P*-value < 0.001 and $\log_2$ fold change > 2) across all comparisons are shown in (A) downregulated DETs are shown in (B) and Upregulated DETs are shown in (C). Pink circle represents 24 hpi treatment vs 24 hpi mock comparison, green circle is 48 hpi treatment vs. 48 hpi mock and yellow is 48 hpi treatment vs 24 hpi treatment.

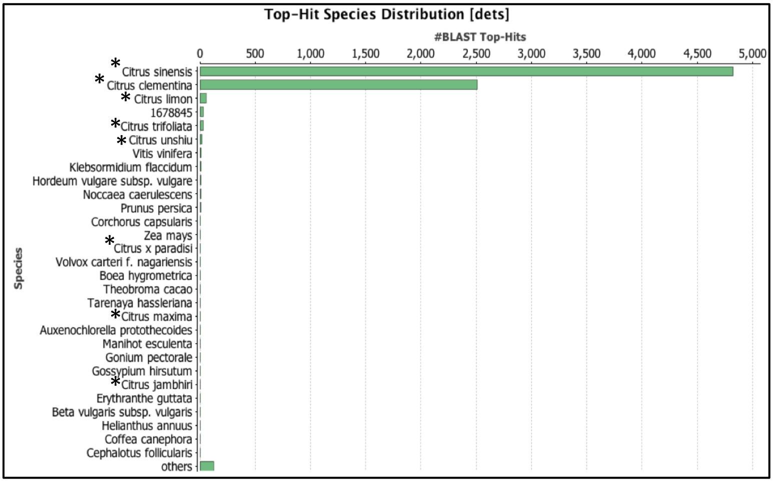


**Figure S5: Top hit species distribution graph.**

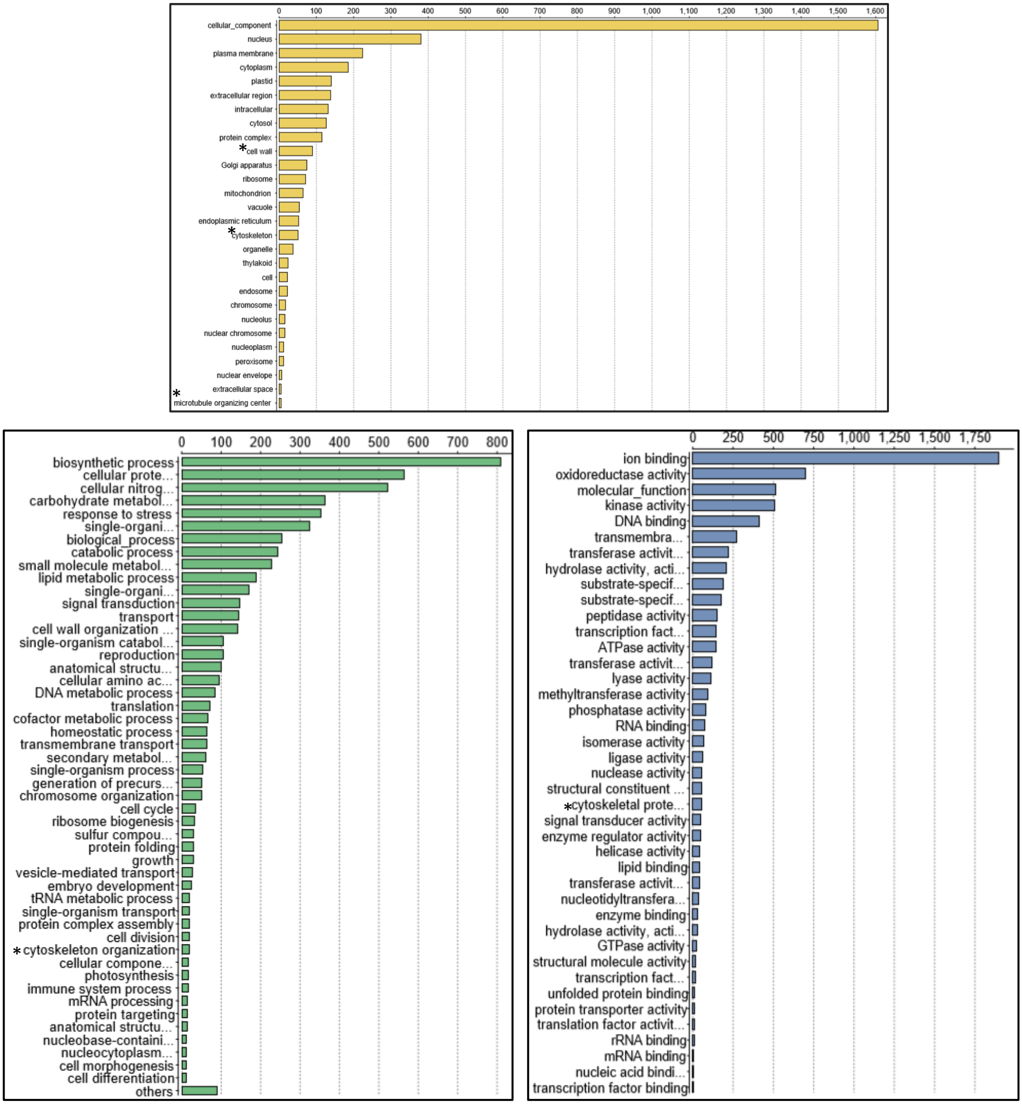


**Figure S6**: Detailed GO term annotations of DETs in biological processes, molecular function and cellular component classes.

## A

| GO Information | | | | CM | | | | ID:545563465 | | ID:814808387 | | ID:312755191 | | ID:456375600 | |
|---|---|---|---|---|---|---|---|---|---|---|---|---|---|---|---|
| No | GO Term | Onto | Description | 1 | 2 | 3 | 4 | Ajusted Pvalue | Num | Ajusted Pvalue | Num | Ajusted Pvalue | Num | Ajusted Pvalue | Num |
| 1 | GO:0004672 | F | protein kinase activity | | | | | 5.9e-42 | 165 | 1.6e-08 | 71 | 9.4e-33 | 161 | 3.3e-14 | 124 |
| 2 | GO:0016773 | F | phosphotransferase activity, alcohol group as acceptor | | | | | 4.6e-35 | 169 | 3e-06 | 73 | 2.7e-32 | 178 | 1.2e-11 | 132 |
| 3 | GO:0005506 | F | iron ion binding | | | | | 1.2e-15 | 42 | 1e-37 | 61 | 3.1e-18 | 49 | 6.6e-47 | 89 |
| 4 | GO:0003824 | F | catalytic activity | | | | | 1.5e-06 | 534 | 1.6e-20 | 460 | 5.7e-10 | 630 | 4.3e-21 | 744 |
| 5 | GO:0004175 | F | endopeptidase activity | | | | | 5.8e-05 | 32 | 0.0036 | 21 | 0.0036 | 30 | 1.7e-07 | 43 |
| 6 | GO:0070001 | F | aspartic-type peptidase activity | | | | | 0.0023 | 15 | 3.7e-05 | 15 | 2.7e-05 | 20 | 1.3e-06 | 23 |
| 7 | GO:0004190 | F | aspartic-type endopeptidase activity | | | | | 0.0023 | 15 | 3.7e-05 | 15 | 2.7e-05 | 20 | 1.3e-06 | 23 |
| 8 | GO:0016209 | F | antioxidant activity | | | | | 0.033 | 17 | 2.9e-07 | 23 | 0.023 | 19 | 1.1e-06 | 30 |
| 9 | GO:0016684 | F | oxidoreductase activity, acting on peroxide as acceptor | | | | | 0.039 | 15 | 3.1e-07 | 21 | 0.047 | 16 | 1.7e-07 | 29 |
| 10 | GO:0004601 | F | peroxidase activity | | | | | 0.039 | 15 | 3.1e-07 | 21 | 0.047 | 16 | 1.7e-07 | 29 |

## B

| GO Information | | | | CM | | | | ID:545563465 | | ID:814808387 | | ID:312755191 | | ID:456375600 | |
|---|---|---|---|---|---|---|---|---|---|---|---|---|---|---|---|
| No | GO Term | Onto | Description | 1 | 2 | 3 | 4 | Ajusted Pvalue | Num | Ajusted Pvalue | Num | Ajusted Pvalue | Num | Ajusted Pvalue | Num |
| 1 | GO:0005488 | F | binding | | | | | 1e-32 | 749 | --- | --- | 3.8e-31 | 833 | 0.0037 | 699 |
| 2 | GO:0016301 | F | kinase activity | | | | | 2.5e-20 | 171 | --- | --- | 5.4e-19 | 184 | 0.00055 | 133 |
| 3 | GO:0005515 | F | protein binding | | | | | 3.2e-16 | 240 | --- | --- | 4.3e-20 | 281 | 0.00055 | 217 |
| 4 | GO:0016772 | F | transferase activity, transferring phosphorus-containing groups | | | | | 6.4e-16 | 174 | --- | --- | 5.4e-15 | 189 | 0.026 | 136 |
| 5 | GO:0016740 | F | transferase activity | | | | | 3.2e-07 | 222 | --- | --- | 3.9e-10 | 264 | 0.0092 | 229 |
| 6 | GO:0004252 | F | serine-type endopeptidase activity | | | | | 0.0015 | 13 | --- | --- | --- | --- | 5e-05 | 17 |
| 7 | GO:0004866 | F | endopeptidase inhibitor activity | | | | | 0.032 | 8 | --- | --- | 0.018 | 9 | --- | --- |
| 8 | GO:0030414 | F | peptidase inhibitor activity | | | | | 0.049 | 8 | --- | --- | 0.03 | 9 | --- | --- |

## C

| GO Information | | | | CM | | | | ID:545563465 | | ID:814808387 | | ID:312755191 | | ID:456375600 | |
|---|---|---|---|---|---|---|---|---|---|---|---|---|---|---|---|
| No | GO Term | Onto | Description | 1 | 2 | 3 | 4 | Ajusted Pvalue | Num | Ajusted Pvalue | Num | Ajusted Pvalue | Num | Ajusted Pvalue | Num |
| 1 | GO:0010876 | P | lipid localization | | | | | --- | --- | 0.0099 | 7 | --- | --- | 6.9e-05 | 12 |
| 2 | GO:0004553 | F | hydrolase activity, hydrolyzing O-glycosyl compounds | | | | | --- | --- | 5.9e-17 | 62 | --- | --- | 4.1e-14 | 80 |
| 3 | GO:0016798 | F | hydrolase activity, acting on glycosyl bonds | | | | | --- | --- | 9.6e-16 | 63 | --- | --- | 1.5e-12 | 81 |
| 4 | GO:0046872 | F | metal ion binding | | | | | --- | --- | 0.00016 | 101 | --- | --- | 0.00024 | 159 |
| 5 | GO:0046914 | F | transition metal ion binding | | | | | --- | --- | 0.00035 | 84 | --- | --- | 0.00085 | 130 |
| 6 | GO:0043169 | F | cation binding | | | | | --- | --- | 0.00065 | 101 | --- | --- | 0.0014 | 159 |
| 7 | GO:0043167 | F | ion binding | | | | | --- | --- | 0.00065 | 101 | --- | --- | 0.0014 | 159 |
| 8 | GO:0016702 | F | oxidoreductase activity, acting on single donors with incorporation of molecular oxygen, incorporation of two atoms of oxygen | | | | | --- | --- | 0.011 | 6 | --- | --- | 0.00013 | 11 |
| 9 | GO:0051213 | F | dioxygenase activity | | | | | --- | --- | 0.017 | 6 | --- | --- | 0.00024 | 11 |
| 10 | GO:0016701 | F | oxidoreductase activity, acting on single donors with incorporation of molecular oxygen | | | | | --- | --- | 0.033 | 6 | --- | --- | 0.00057 | 11 |
| 11 | GO:0003774 | F | motor activity | | | | | --- | --- | 0.041 | 10 | --- | --- | 0.00044 | 19 |

## D

| GO Information | | | | CM | | | | ID:545563465 | | ID:814808387 | | ID:312755191 | | ID:456375600 | |
|---|---|---|---|---|---|---|---|---|---|---|---|---|---|---|---|
| No | GO Term | Onto | Description | 1 | 2 | 3 | 4 | Ajusted Pvalue | Num | Ajusted Pvalue | Num | Ajusted Pvalue | Num | Ajusted Pvalue | Num |
| 1 | GO:0003777 | F | microtubule motor activity | | | | | --- | --- | --- | --- | --- | --- | 8.9e-05 | 18 |
| 2 | GO:0017171 | F | serine hydrolase activity | | | | | --- | --- | --- | --- | --- | --- | 0.00024 | 28 |
| 3 | GO:0008236 | F | serine-type peptidase activity | | | | | --- | --- | --- | --- | --- | --- | 0.00024 | 28 |
| 4 | GO:0046983 | F | protein dimerization activity | | | | | --- | --- | --- | --- | --- | --- | 0.00055 | 23 |
| 5 | GO:0070011 | F | peptidase activity, acting on L-amino acid peptides | | | | | --- | --- | --- | --- | --- | --- | 0.0021 | 56 |
| 6 | GO:0000287 | F | magnesium ion binding | | | | | --- | --- | --- | --- | --- | --- | 0.012 | 12 |
| 7 | GO:0016747 | F | transferase activity, transferring acyl groups other than amino-acyl groups | | | | | --- | --- | --- | --- | --- | --- | 0.014 | 31 |
| 8 | GO:0008233 | F | peptidase activity | | | | | --- | --- | --- | --- | --- | --- | 0.021 | 56 |
| 9 | GO:0016746 | F | transferase activity, transferring acyl groups | | | | | --- | --- | --- | --- | --- | --- | 0.034 | 32 |
| 10 | GO:0004180 | F | carboxypeptidase activity | | | | | --- | --- | --- | --- | --- | --- | 0.038 | 12 |
| 11 | GO:0005507 | F | copper ion binding | | | | | --- | --- | --- | --- | --- | --- | 0.043 | 15 |
| 12 | GO:0016021 | C | integral to membrane | | | | | --- | --- | --- | --- | --- | --- | 6.1e-09 | 60 |

**Figure S7. Snap shot of SEACOMPARE.** It is Cross comparison of Singular Enrichment Analysis (SEA).

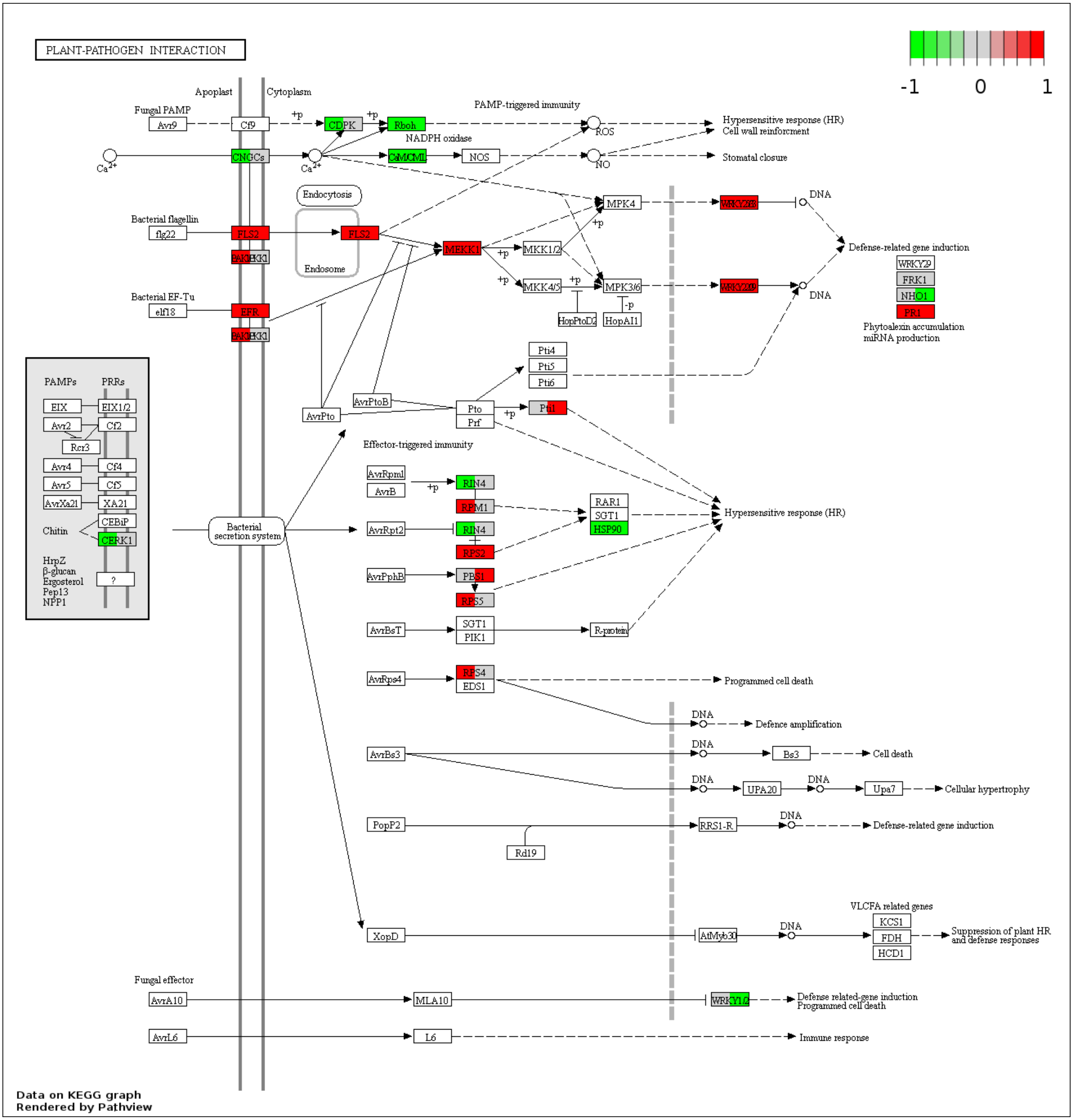

PLANT-PATHOGEN INTERACTION
-1
0
1
Apoplast
Cytoplasm
Fungal PAMP
Avr9
Cf9
+p
CDPK
Rboh
NADPH oxidase
PAMP-triggered immunity
ROS
Hypersensitive response (HR)
Cell wall reinforcment
Ca2+
CNGCs
CaM/CML
NOS
NO
Stomatal closure
Endocytosis
Endosome
Bacterial flagellin
flg22
FLS2
BAK1
BKK1
MEKK1
MKK1/2
MPK4
WRKY25/33
DNA
Defense-related gene induction
WRKY29
FRK1
NHO1
PR1
Phytoalexin accumulation
miRNA production
MKK4/5
MPK3/6
WRKY22/29
HopPtoD2
HopAI1
-p
Bacterial EF-Tu
elf18
EFR
PAMPs
PRRs
EIX
EIX1/2
Avr2
Cf2
Rcr3
Avr4
Cf4
Avr5
Cf5
AvrXa21
XA21
CEBiP
Chitin
CERK1
HrpZ
β-glucan
Ergosterol
Pep13
NPP1
?
Pti4
Pti5
Pti6
AvrPtoB
AvrPto
Pto
Prf
Pti1
Effector-triggered immunity
AvrRpm1
AvrB
RIN4
RPM1
RAR1
SGT1
HSP90
Hypersensitive response (HR)
Bacterial secretion system
AvrRpt2
RPS2
AvrPphB
PBS1
RPS5
AvrBsT
PIK1
R-protein
AvrRps4
RPS4
EDS1
Programmed cell death
Defence amplification
AvrBs3
Bs3
Cell death
UPA20
Upa7
Cellular hypertrophy
PopP2
RRS1-R
Rd19
VLCFA related genes
KCS1
FDH
HCD1
XopD
AtMyb30
Suppression of plant HR and defense responses
Fungal effector
AvrA10
MLA10
WRKY1/2
Defense related-gene induction
AvrL6
L6
Immune response
Data on KEGG graph
Rendered by Pathview

**Figure S8. Mapping of differentially expressed transcripts on Plant-pathogen interaction pathway.** Color gradient represents log2 fold ratios with red representing upregulation and green representing downregulation in treatments over mock roots. Left and right-hand side of each KO box show DETs mapping at 24hpi and 48hpi respectively. White box or no color fill means no DET was assigned to that KO term. Half white box on either side means that term was not differentially expressed at the respective time point.

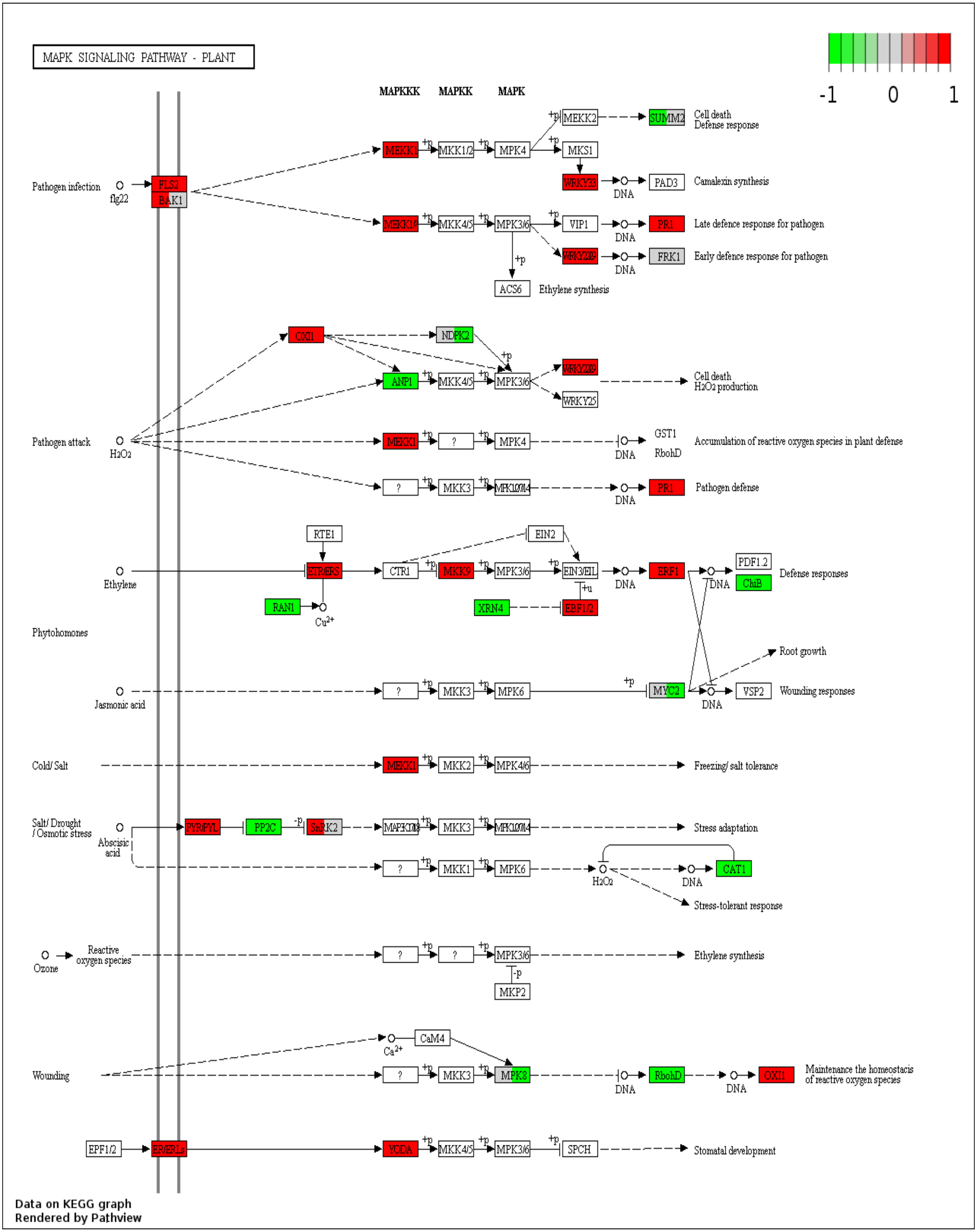
MAPK SIGNALING PATHWAY - PLANT
-1
0
1
MAPKKK
MAPKK
MAPK
Pathogen infection
flg22
FLS2
BAK1
MEKK1
MKK1/2
MPK4
MEKK2
SUMM2
Cell death
Defense response
MKS1
WRKY33
DNA
PAD3
Camalexin synthesis
MEKK1/
MKK4/5
MPK3/6
VIP1
PR1
Late defence response for pathogen
WRKY22/29
FRK1
Early defence response for pathogen
ACS6
Ethylene synthesis
OXI1
NDPK2
ANP1
WRKY25
Cell death
H2O2 production
Pathogen attack
H2O2
GST1
RbohD
Accumulation of reactive oxygen species in plant defense
MKK3
MPK1/2/7/14
Pathogen defense
RTE1
EIN2
Ethylene
ETR/ERS
CTR1
MKK9
EIN3/EIL
ERF1
PDF1.2
ChiB
Defense responses
RAN1
Cu2+
XRN4
EBF1/2
Phytohomones
Root growth
Jasmonic acid
MPK6
MYC2
VSP2
Wounding responses
Cold/ Salt
MKK2
MPK4/6
Freezing/ salt tolerance
Salt/ Drought
/ Osmotic stress
Abscisic acid
PYR/PYL
PP2C
SnRK2
MAPKKK17/18
Stress adaptation
MKK1
CAT1
Stress-tolerant response
Ozone
Reactive oxygen species
MKP2
Ca2+
CaM4
Wounding
MPK8
Maintenance the homeostacis of reactive oxygen species
EPF1/2
ER/ERLs
YODA
SPCH
Stomatal development
Data on KEGG graph
Rendered by Pathview

**Figure S9. Visualization of differentially expressed transcripts involved in MAPK signaling pathway.** Color gradient represents log2 fold ratios with red representing upregulation and green representing downregulation in treatments over mock roots. Left and right-hand side of each KO box show DETs mapping at 24hpi and 48hpi respectively. White box or no color fill means no DET was assigned to that KO term. Half white box on either side means that term was not differentially expressed at the respective time point. Significant dominance of up regulated transcripts was seen in whole pathway.

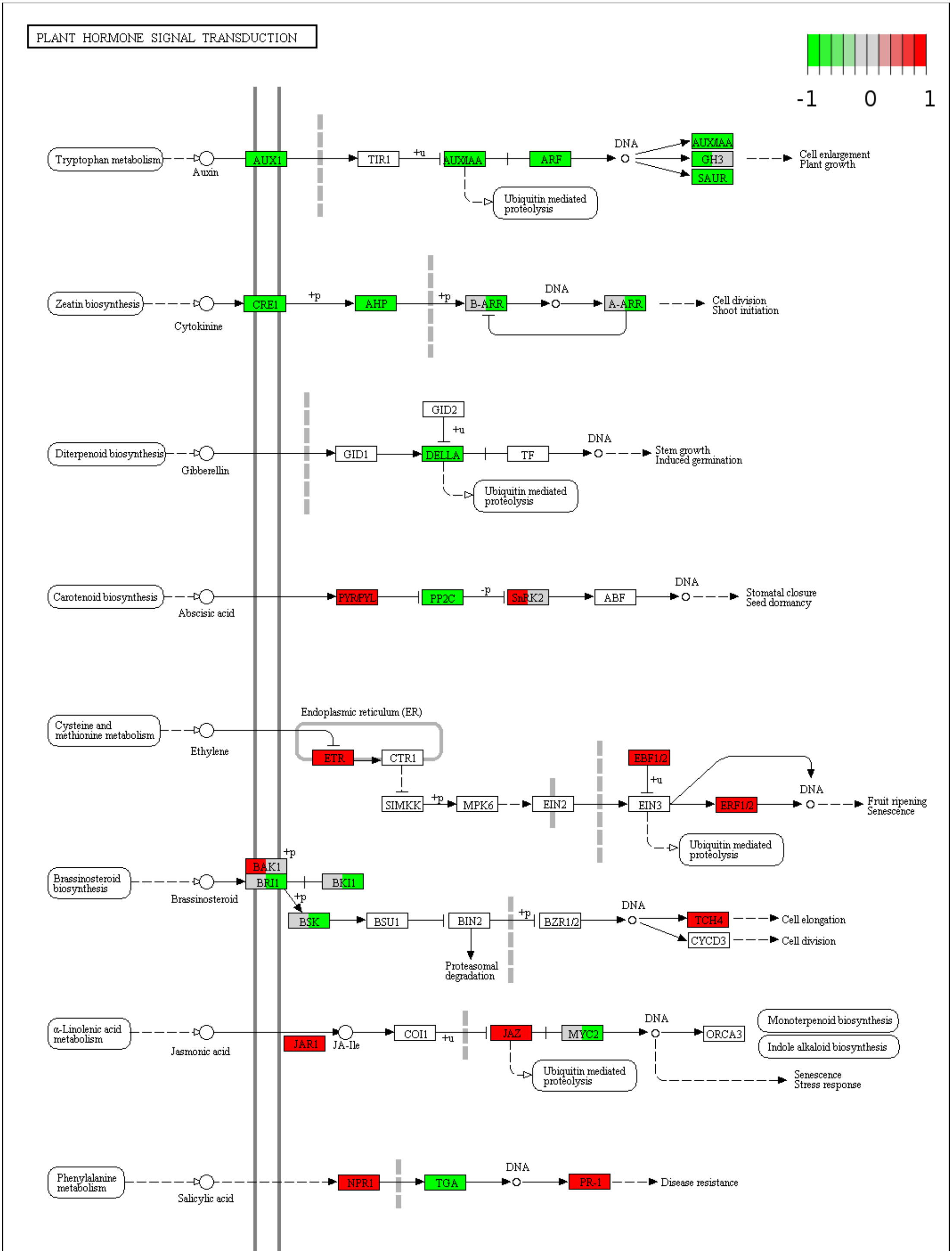


Data on KEGG graph
Rendered by Pathview

**Figure S10. Differentially expressed transcripts mapped on Plant hormone signaling pathway.** Color gradient represents log2 fold ratios with red representing upregulation and green representing downregulation in treatments over mock roots. Left and right-hand side of each KO box show DETs mapping at 24hpi and 48hpi respectively. White box or no color fill means no DET was assigned to that KO term. Half white box on either side means that term was not differentially expressed at the respective time point.

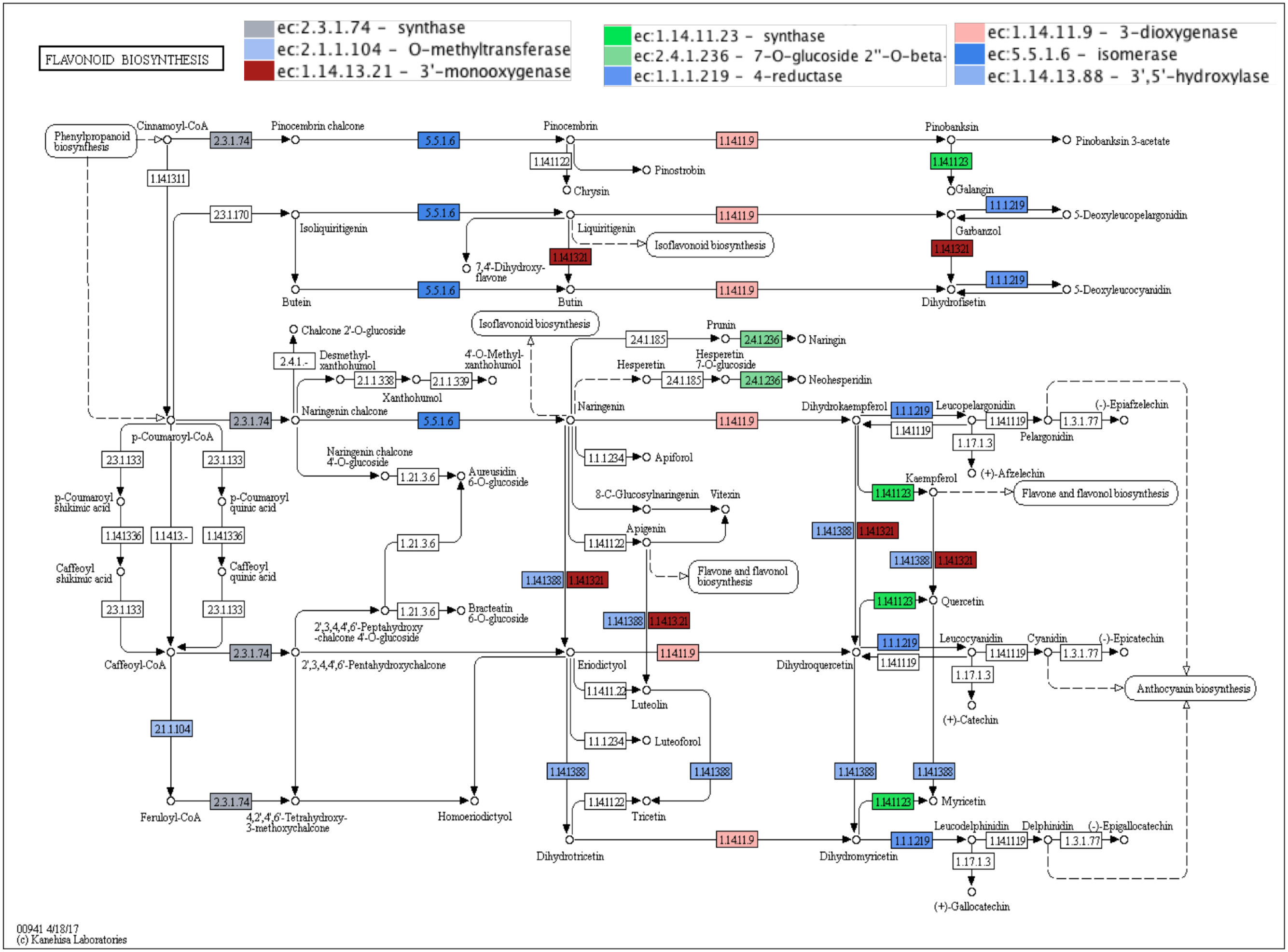
FLAVONOID BIOSYNTHESIS
ec:2.3.1.74 - synthase
ec:2.1.1.104 - O-methyltransferase
ec:1.14.13.21 - 3'-monooxygenase
ec:1.14.11.23 - synthase
ec:2.4.1.236 - 7-O-glucoside 2''-O-beta-
ec:1.1.1.219 - 4-reductase
ec:1.14.11.9 - 3-dioxygenase
ec:5.5.1.6 - isomerase
ec:1.14.13.88 - 3',5'-hydroxylase
Phenylpropanoid biosynthesis
Cinnamoyl-CoA
Pinocembrin chalcone
Pinocembrin
Pinobanksin
Pinobanksin 3-acetate
Pinostrobin
Chrysin
Galangin
5-Deoxyleucopelargonidin
Isoliquiritigenin
Liquiritigenin
Isoflavonoid biosynthesis
7,4'-Dihydroxy-flavone
Garbanzol
Butein
Butin
Dihydrofisetin
5-Deoxyleucocyanidin
Chalcone 2'-O-glucoside
Desmethyl-xanthohumol
4'-O-Methyl-xanthohumol
Xanthohumol
Prunin
Naringin
Hesperetin
Hesperetin 7-O-glucoside
Neohesperidin
p-Coumaroyl-CoA
Naringenin chalcone
Naringenin
Dihydrokaempferol
Leucopelargonidin
Pelargonidin
(-)-Epiafzelechin
(+)-Afzelechin
Naringenin chalcone 4'-O-glucoside
Aureusidin 6-O-glucoside
Apiforol
8-C-Glucosylnaringenin
Vitexin
Apigenin
Kaempferol
Flavone and flavonol biosynthesis
p-Coumaroyl shikimic acid
p-Coumaroyl quinic acid
Caffeoyl shikimic acid
Caffeoyl quinic acid
2',3,4,4',6'-Peptahydroxy-chalcone 4'-O-glucoside
Bracteatin 6-O-glucoside
Quercetin
Caffeoyl-CoA
2',3,4,4',6'-Pentahydroxychalcone
Eriodictyol
Dihydroquercetin
Leucocyanidin
Cyanidin
(-)-Epicatechin
(+)-Catechin
Anthocyanin biosynthesis
Luteolin
Luteoforol
Feruloyl-CoA
4,2',4',6'-Tetrahydroxy-3-methoxychalcone
Homoeriodictyol
Tricetin
Myricetin
Dihydrotricetin
Dihydromyricetin
Leucodelphinidin
Delphinidin
(-)-Epigallocatechin
(+)-Gallocatechin
00941 4/18/17
(c) Kanehisa Laboratories

**Figure S11. KEGG enzyme enrichment analysis showed maximum mapping on Flavanoid biosynthesis pathway.** Color key represents names of different enzymes involved in the pathway.

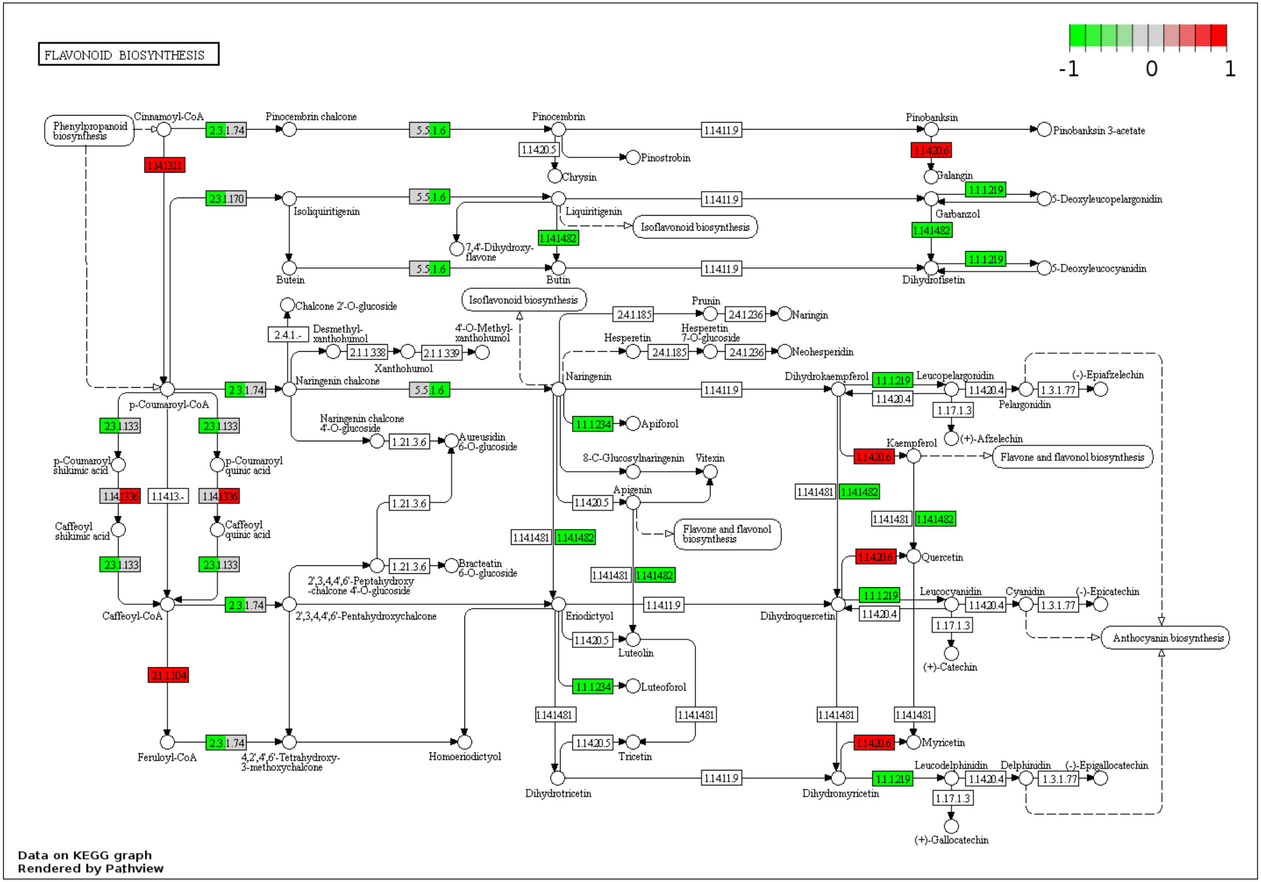

FLAVONOID BIOSYNTHESIS
-1
0
1
Phenylpropanoid biosynthesis
Cinnamoyl-CoA
Pinocembrin chalcone
Pinocembrin
Pinobanksin
Pinobanksin 3-acetate
Pinostrobin
Chrysin
Galangin
Isoliquiritigenin
Liquiritigenin
Isoflavonoid biosynthesis
Garbanzol
5-Deoxyleucopelargonidin
7,4'-Dihydroxy-flavone
Butein
Butin
Dihydrofisetin
5-Deoxyleucocyanidin
Chalcone 2'-O-glucoside
Prunin
Naringin
Desmethyl-xanthohumol
4'-O-Methyl-xanthohumol
Xanthohumol
Hesperetin
Hesperetin 7-O-glucoside
Neohesperidin
Naringenin chalcone
Naringenin
Dihydrokaempferol
Leucopelargonidin
Pelargonidin
(-)-Epiafzelechin
(+)-Afzelechin
p-Coumaroyl-CoA
Apiforol
Naringenin chalcone 4'-O-glucoside
Aureusidin 6-O-glucoside
Kaempferol
Flavone and flavonol biosynthesis
p-Coumaroyl shikimic acid
p-Coumaroyl quinic acid
8-C-Glucosylnaringenin
Vitexin
Apigenin
Caffeoyl shikimic acid
Caffeoyl quinic acid
Quercetin
2',3,4,4',6'-Peptahydroxy-chalcone 4'-O-glucoside
Bracteatin 6-O-glucoside
Caffeoyl-CoA
2',3,4,4',6'-Pentahydroxychalcone
Eriodictyol
Dihydroquercetin
Leucocyanidin
Cyanidin
(-)-Epicatechin
Anthocyanin biosynthesis
Luteolin
(+)-Catechin
Luteoforol
Feruloyl-CoA
4,2',4',6'-Tetrahydroxy-3-methoxychalcone
Homoeriodictyol
Tricetin
Myricetin
Dihydrotricetin
Dihydromyricetin
Leucodelphinidin
Delphinidin
(-)-Epigallocatechin
(+)-Gallocatechin
2.3.1.74
5.5.1.6
1.14.11.9
1.14.20.5
1.14.20.6
1.14.13.11
2.3.1.170
1.1.1.219
1.14.14.82
2.4.1.-
2.4.1.185
2.4.1.236
2.1.1.338
2.1.1.339
1.14.20.4
1.3.1.77
1.17.1.3
2.3.1.133
1.1.1.234
1.21.3.6
1.14.13.36
1.14.13.-
1.14.14.81
2.1.1.104
Data on KEGG graph
Rendered by Pathview

**Figure S12. KEGG enzyme enrichment analysis showed maximum mapping on Flavanoid biosynthesis pathway.** Color key represents names of different enzymes involved in the pathway.
